# Evidence for complex interplay between quorum sensing and antibiotic resistance in *Pseudomonas aeruginosa*

**DOI:** 10.1101/2022.04.05.487235

**Authors:** Rakesh Sikdar, Mikael H. Elias

## Abstract

<u>Q</u>uorum <u>s</u>ensing (QS) is a cell-density-dependent, intercellular communication system mediated by small diffusible signaling molecules. QS regulates a range of bacterial behaviors, including biofilm formation, virulence, drug resistance mechanisms, and antibiotic tolerance. Enzymes capable of degrading signaling molecules can interfere in QS - a process termed <u>q</u>uorum <u>q</u>uenching (QQ). Remarkably, previous work reported some cases where enzymatic interference in QS was synergistic to antibiotics against *Pseudomonas aeruginosa*. The premise of combination therapy is attractive to fight against multidrug-resistant bacteria, yet comprehensive studies are lacking. Here we evaluate the effects of QS signal disruption on the antibiotic resistance profile of *P. aeruginosa* by testing 222 antibiotics and antibacterial compounds from 15 different classes. We found compelling evidence that QS signal disruption does indeed affect antibiotic resistance (40% of all tested compounds; 89/222), albeit not always synergistically (not synergistic for 48% of compounds with an effect (43/89)). For some tested antibiotics, like sulfathiazole and trimethoprim, we were able to relate the changes in resistance caused by QS signal disruption to the modulation of the expression of key genes of the folate biosynthetic pathway. Moreover, using a *P. aeruginosa*-based *Caenorhabditis elegans* killing model, we confirm that enzymatic QQ modulates the effects of antibiotics on *P. aeruginosa’s* pathogenicity *in vivo*. Altogether, these results show that signal disruption has profound and complex effects on the antibiotic resistance profile of *P. aeruginosa*. This work suggests that combination therapy including QQ and antibiotics should not be discussed globally but rather in case-by-case studies.

**IMPORTANCE:** Interference in bacterial Quorum Sensing (QS) is a promising approach to control microbial behavior. Of particular interest is the potential of this strategy to reduce biofilms and virulence of antibiotic resistant strains. Interestingly, several studies report synergistic interactions between antibiotic treatments and interference in QS. However, it is unclear whether this is a generality, let alone the molecular mechanisms underlying the observed synergies. Here, we provide a comprehensive description of combination treatment in the model organism, opportunistic human pathogen *P. aeruginosa*. Screening > 200 antimicrobials, and combining them to QS signals and disruption strategies, we show that there is no systematic synergy between these approaches *in vitro*, as well as *in vivo*, in a *C. elegans* infection model. Altogether, this work show that QS has complex connections to the antibiotic resistance profile of *P. aeruginosa*, and that combination treatment should not be discussed globally, but rather in case-by-case studies.

## INTRODUCTION

Antibiotic resistance is reported to be rapidly rising, possibly partly due to the overuse of antibiotics in medical, agricultural, and industrial applications [1–3]. This risk may have increased during the COVID-19 pandemic and the increased use of antibiotics to prevent secondary bacterial infections in hospitals [4]. Pathogenic bacteria that are relevant in human diseases, such as *Pseudomonas aeruginosa,* can be resistant to numerous antibiotic treatments [5]. It is associated with 10% of nosocomial infections [6] and is the main cause of mortality and morbidity in a debilitating genetic disease like Cystic Fibrosis in humans [7]. It is listed among the top priority pathogens by the WHO for immediate R&D of new antimicrobials [8]. *P. aeruginosa* shows a remarkable ability to adapt to a wide range of environmental niches due to its high genome plasticity [9, 10]. Interestingly, the virulence of *P. aeruginosa*, like many other pathogenic microbes, is regulated by a chemical communication system termed <u>Q</u>uorum <u>S</u>ensing (QS) [11]. Consequently, interference in QS signaling is appealing to control microbial pathogens.

Numerous bacteria use QS for communication: they produce, secrete, sense, and respond to small diffusible signaling molecules known as <u>A</u>utoInducers (AIs). One main class of autoinducers is autoinducer-I or *N-*<u>A</u>cyl <u>H</u>omoserine <u>L</u>actones (AHLs). AHL-based QS circuits are reported to regulate the expression of up to 6% of bacterial genes [12] and to modulate bacterial behaviors critical for their pathogenicity, such as virulence factor production, drug resistance, toxin production, motility, and biofilm formation, in a cell density-dependent manner [13].

*P. aeruginosa* has three interwoven QS signaling circuits with overlapping genetic targets - namely LasIR, RhlIR, and *Pseudomonas* Quinolone Signal (PQS) in a top-to-bottom order of hierarchy [14, 15]. They produce, detect, and respond to autoinducer molecules *N*-3- oxo-dodecanoyl-*L*-Homoserine Lactone (3oC12-HSL), *N*-butyryl-*L*-Homoserine lactone (C4-HSL) and alkyl-quinolones respectively. This sophisticated QS circuitry enables this bacterium to be a versatile and opportunistic pathogen that can adapt to a variety of environmental conditions in the host tissue and forms a robust biofilm that is difficult to disperse [16, 17].

The antibiotic resistance of *P. aeruginosa* stems from several intrinsic, acquired, and adaptive mechanisms – as elaborated in the following Reviews [5,18,19]. These mechanisms can be three-fold - (i) Chemical modification of antibiotics using enzymes such as β–lactamases [20], aminoglycoside modifying enzymes [21], 16s rRNA methylases [22]; (ii) Modification of biofilm structure (extracellular polymeric substances) [23], membrane physiology (outer membrane permeability, LPS modification) [24] and/or surface porins (OprF, OprD, and OprH) to reduce antibiotic permeability [25]; (iii) Expression of multidrug efflux pumps (MexAB-OprM, MexCD-OprJ, MexEF-OprN, and MexXY-OprM) to secrete the antibiotics out of the cell [19], (iv) Altering the expression and/or characteristics of genes and proteins targeted by antibiotics (e.g. DNA gyrases, folate biosynthetic pathway genes) [22] and (v) utilizing global stress response systems (two-component signaling systems, e.g. PhoPQ, CprRS, ParRS) and phenotypic modifications (swarming and surfing motility, LPS modifications) to adapt to the antibiotic-mediated stress [26–28]. Many of these mechanisms may be regulated by QS regulation [18, 19] and therefore antibiotic resistance and QS are possibly interconnected. In support of this hypothesis, some previous studies have highlighted potential synergistic effects between antibiotic treatments and interference in QS [29–36] yet a comprehensive investigation is lacking.

Numerous enzymes capable of hydrolyzing AHLs were isolated and characterized [13, 37]. Lactonases, enzymes that hydrolyze and open the lactone ring of AHLs, have been enzymatically and structurally well-studied [38, 39]. Well-characterized representatives from the Phosphotriesterase-Like Lactonase (PLL) family include VmoLac [40], SisLac [41], PPH [42] or SsoPox [43–45], as well as representatives from the Metallo-β-lactamase Lactonase (MLL) such as MomL [46], AiiA [47], AaL [48] or GcL [49]. By hydrolyzing AHLs, lactonases interfere in QS, a process termed <u>Q</u>uorum <u>Q</u>uenching (QQ). QQ enzymes were shown to reduce the virulence of *P. aeruginosa* both *in vitro* and *in vivo* [46,50–58]. As *P. aeruginosa* utilizes C4-HSL and 3- oxo-C12 HSL-based QS signaling circuits, the AHL preference of the QQ enzyme is important as it may quench either or both LasIR and RhlIR QS circuits. Moreover, recent studies showed that the substrate specificity of the QQ enzyme affected proteome profiles, virulence factors expression, virulence, and biofilm formation in *P. aeruginosa* [56, 59].

In this study, we used two QQ lactonases with distinct substrate specificity, SsoPox [43] and GcL [49], to evaluate the effects of AHL signal disruption on the antibiotic-resistance profile of *P. aeruginosa*. We observed that signal disruption has complex effects on antibiotic resistance that are dependent on the antibiotic/antimicrobial compound and the QQ enzyme used. We confirmed key observations in independent assays. For the antibiotics sulfathiazole and trimethoprim, we provide evidence that changes in key genes regulation due to interference in QS signaling are responsible for the observed modulation of antibiotic resistance using <u>q</u>uantitative <u>R</u>everse <u>T</u>ranscription <u>P</u>olymerase <u>C</u>hain <u>R</u>eaction (qRT-PCR). Lastly, we demonstrate that these effects on antibiotic sensitivity of *P. aeruginosa* can translate *in vivo*, in a *Caenorhabditis elegans* killing assay.

## Materials and Methods

### Reagents, *Pseudomonas* strains, and growth conditions

*Pseudomonas aeruginosa* strain UCBPP-PA14 [60], designated as PA14 throughout this article, is used in all experiments and maintained in the laboratory following standardized protocols [61]. PA14 was cultured in either Miller’s Luria-Bertani (LB) broth (BD Difco, #244610), LB-agar plates (BD Difco, #244510), or in a peptone-based proprietary growth media – GN IF-10A inoculating fluid (Biolog, #72264). Unless otherwise mentioned, all cultures were grown at 37°C with liquid cultures shaking at 250 rpm. All phenotype microarray plates – PM11C (#12211), PM12B (#12212), PM13B (#12213), PM14A (#12214), PM15B (#12215), PM16A (#12216), PM17A (#12217), PM18C (#12218), PM19 (#12219) and PM20B (#12220) and 100x Dye Mix A (#74221) were purchased from Biolog (Hayward, CA). *N-*butyryl-*L*-Homoserine lactone designated as C4-HSL (#10007898) and *N*-3-oxo-dodecanoyl-*L*-Homoserine lactone designated as 3oC12 HSL (#10007895) were purchased from Cayman Chemical Company (Ann Arbor, MI) and dissolved in 100% DMSO just before use. All other chemicals including antibiotics were of at least reagent grade and were purchased from either Millipore Sigma (Burlington, MA) or Fisher Scientific (Hampton, NH).

### Lactonase production, purification, and quantitation

Production of the inactive SsoPox mutant 5A8 [62], SsoPox W263I [45] and GcL [49] in *Escherichia coli* strain BL21(DE3) was performed as previously described [44,49,59,63]. All lactonase preparations used in this study were made in buffer (PTE) composed of 50 mM HEPES pH 8.0, 150 mM NaCl, 0.2 mM CoCl_2_, sterilized by passing through a 0.2μ filter, and stored at 4°C until use.

### Preparation of antibiotic stocks

Antibiotic stocks were prepared just before use by dissolution in deionized water (Nafcillin sodium, Azlocillin sodium, Colistin sulfate, Sulfathiazole sodium, Sulfadiazine sodium, D-cyclo-serine, Carbenicillin disodium, and Procaine hydrochloride), in 100% DMSO (Oxacillin sodium, Coumarin, Trimethoprim, and Carbonyl cyanide 3-chlorophenylhydrazone), in 0.1N HCl (Norfloxacin) or 1N NaOH (Ofloxacin). Antibiotic stocks were sterilized by passing through a 0.2μ filter.

### Biolog Phenotype MicroArray^TM^ experiments

These experiments were conducted using a modification of a standard protocol obtained from Biolog Inc (Dr. Barry Bochner, direct communication). Phenotype MicroArray^TM^ (PM) microplates allow for the testing of a range of antibiotics, inorganic salts, and bacteriostatic/bactericidal compounds (10 plates, labeled PM11 through PM20: each containing 24 compounds). Each tested compound is present at 4 concentrations, yet the precise concentration value is not disclosed by Biolog. Therefore, these MicroArrays provide a qualitative estimate of microbial sensitivity. The system is designed to use a proprietary peptone-based growth media – GN IF-10A to be supplemented with a proprietary tetrazolium redox dye mix. The dye is reduced to purple-colored formazan products λabs 590 nm) due to NADH production (a sensitive indicator of respiration) by metabolically active cells [64, 65]. This colorimetric reaction is monitored and recorded by a spectrophotometer at specific time intervals to generate a kinetic response curve mirroring microbial growth that can be further quantitated into parameters such as lag, slope, and area under the curve [66].

A single colony of PA14, picked from freshly streaked LB-agar plates, was inoculated in 2 μmL LB media containing appropriate treatments (50 μg/mL lactonases or *N*-acyl homoserine lactones at a final concentration of 10 μM) and grown at 37°C/250 rpm until cells reach early log phase of growth (OD_600_ ∼ 0.2 - 0.3). Cultures were then harvested by centrifugation (5000g for 3 min) and washed three times with Biolog IF-10A media (1.2x GN IF- 10A inoculating fluid diluted to 1x using autoclaved deionized water and containing 1x Biolog dye mix A). Washed cells were resuspended in Biolog IF-10A media at an OD_750_ of ∼ 0.05. A final inoculum solution is made by diluting the washed cells 1:100 into fresh Biolog IF-10μA media containing the similar treatment as above (50 μg/mL lactonases or *N*-acyl homoserine lactones at a final concentration of 10 μL of the final inoculum solution was dispensed into each well of a 96 well Phenotype MicroArray microplate and overlayed with 20 μL sterile light mineral oil (Fujifilm Irvine #9305) to limit evaporation. Antibiotic-free Growth of PA14 with 50 μg/mL SsoPox 5A8 for control purposes was carried out on a half-area sterile 96-well microplate (Corning #3696) to mimic the physical dimensions of Biolog PM microplates, using a similar protocol as above. The microplate was then incubated with a lid in a BioTek Epoch2 microplate spectrophotometer at 37°C with linear shaking at 300 rpm for 24 hrs. OD_590_ (absorbance of formazan, as discussed earlier) and OD_750_ (for cell turbidity) were measured for each well every 15 min. The [OD_590_ – OD_750_] values plotted against the time of growth corresponds to the growth curve of PA14 and the area-under-the-curve (AUC), calculated with the BioTek Gen5 (v2.9) software corresponds to the total growth of PA14 over 24 h.

### Data analysis and presentation for the Biolog Phenotype MicroArray experiments

Raw area-under-the-curve (AUC) data from the BioTek Gen5 software was analyzed and processed initially in Microsoft Excel worksheets before final analysis in GraphPad Prism 8. AUC values less than 1000 were found to be baseline noise and were discarded from subsequent analysis. Each antibiotic or antibacterial compound in the MicroArrays has 4 gradually increasing dosages (undisclosed by Biolog) in 4 horizontally adjacent microwells (denoted as “Concentration 1” through “Concentration 4” in **Fig. S1A, B - S5A, B**, with the leftmost microwell containing the lowest dose (Concentration 1) and the rightmost microwell containing the highest dose (Concentration 4). The AUC data was used to compare the treatments by normalizing each treatment (SsoPox W263I, GcL, C4-HSL, 3oC12-HSL) to the control (SsoPox 5A8; inactive lactonase) using the following formula:

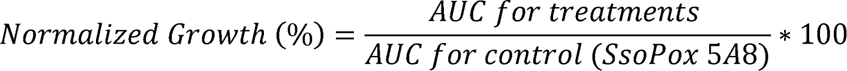

We present the treatment-dependent normalized results for all the 222 unique antibiotics and antibacterial compounds, organized according to their dosage (Concentration 1-4) in the form of graded heatmaps (white-red-black color schema) in **Fig. S1-S5.** An additional heatmap strip with a green-black color scheme is overlayed on the right, which represents the sensitivity of PA14 to the indicated concentration of the compounds. We term it as the “relative resistance score” of PA14 against that compound at that indicated concentration, calculated by the following formula:

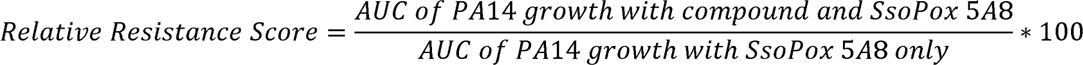

A relative resistance score of 100% means that PA14 is completely resistant to the tested compound and that its growth is unaffected by the compound. All heatmaps were generated and processed in GraphPad Prism 8. Shorter synonyms of the following compound names used by Biolog in the MicroArrays were obtained from PubChem (https://pubchem.ncbi.nlm.nih.gov/) and used in the heat maps (**Fig. 1, 2, and S7**) for aesthetic purposes and indicated in braces - Methyltrioctylammoniumchloride (Methyltrioctyl-NH4Cl); 5,7-Dichloro-8-hydroxyquinoline (Chloroxine), Sodium pyrophosphate decahydrate (Sodium pyrophosphate); 1-Chloro-2,4- dinitrobenzene (Chlorodinitrobenzene); 5-Nitro-2-furaldehyde semicarbazone (Nitrofurazone); 1- Hydroxypyridine-2-thione (Pyrithione); 3, 4-Dimethoxybenzyl alcohol (Veratrole alcohol) and L-Glutamic-g-hydroxamate (Glutamine hydroxamate).

**Figure 1.**
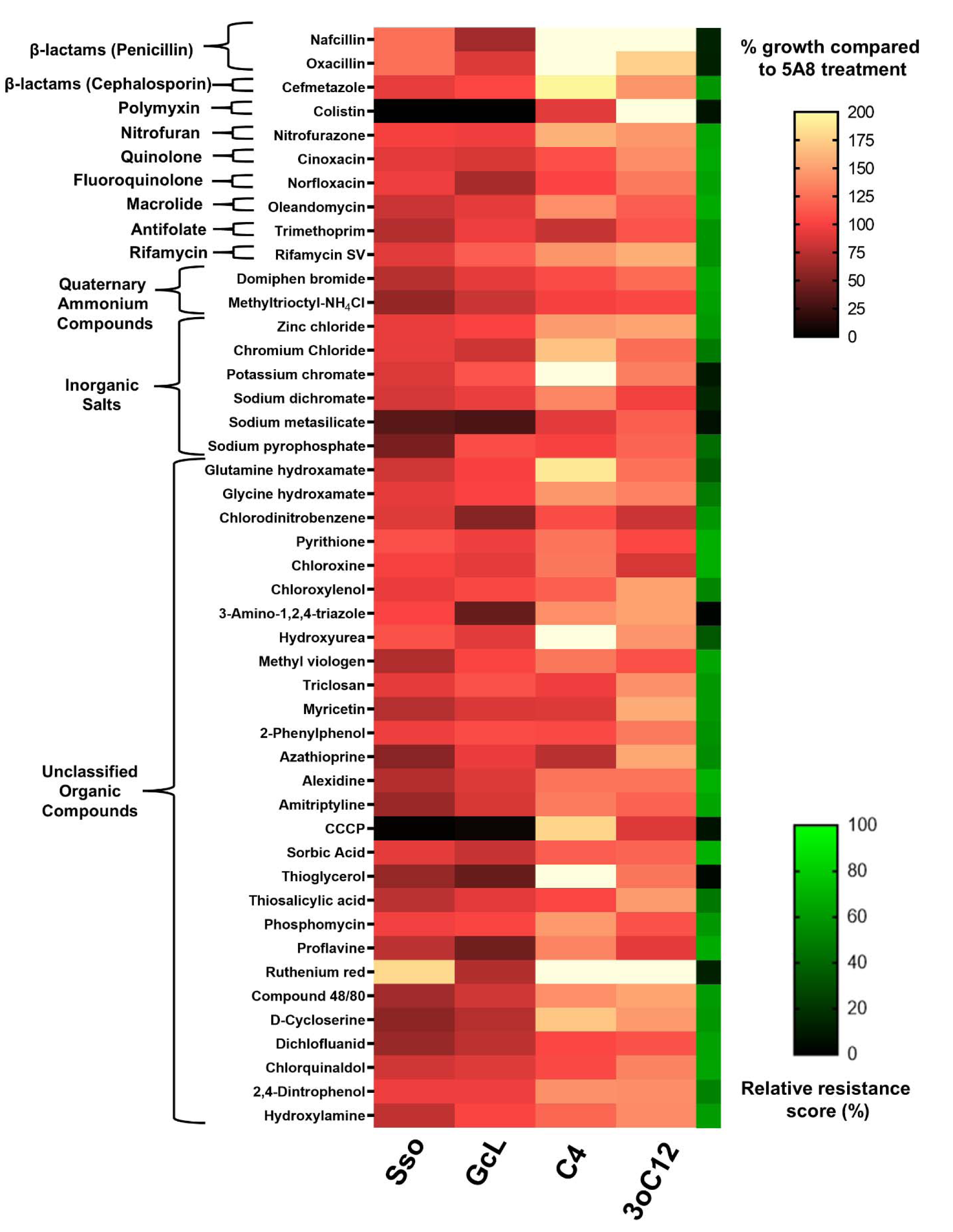
PA14 growth profile for antimicrobials for which lactonase treatment increases sensitivity. Growth of PA14 in the presence of lactonases – SsoPox W263I (Sso) and GcL or exogenously added AHLs – C4-HSL (C4) and 3-oxo-C12-HSL (3oC12) is represented as a % of PA14 growth in the presence of inactive lactonase SsoPox 5A8 (control) with a white-red-black color scheme. Adjacent to it, an additional heat map strip with a green-black color scheme is overlaid, representing the relative resistance score (see Methods) on a % scale, and indicating the sensitivity of PA14 to the tested antibiotics and antibacterial compounds. Tested compounds are grouped according to their classes indicated on the left. Higher relative resistance scores indicate higher resistance of PA14. All lactonases and AHLs are used at 50 μ M final concentrations, respectively.

**Figure 2.**
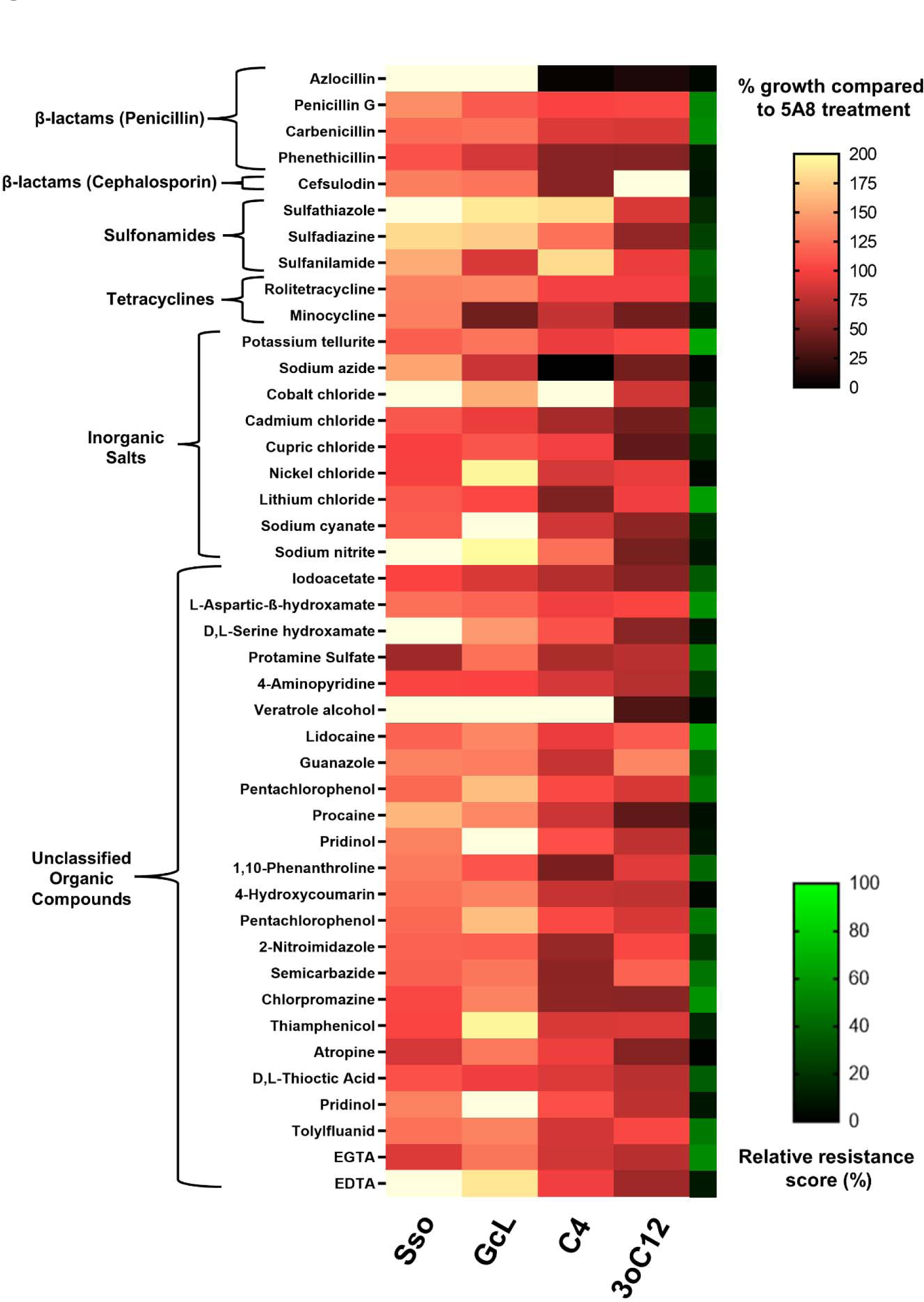
PA14 growth profile for antimicrobials for which lactonase treatment increases resistance. Growth of PA14 in the presence of lactonases – SsoPox W263I (Sso) and GcL or exogenously added AHLs – C4-HSL (C4) and 3-oxo-C12-HSL (3oC12) is represented as a % of PA14 growth in the presence of inactive lactonase SsoPox 5A8 (control) with a white-red-black color scheme. Adjacent to it, an additional heat map strip with a green-black color scheme is overlaid, representing the relative resistance score (see Methods) on a % scale, and indicating the sensitivity of PA14 to the tested antibiotics and antibacterial compounds. Tested compounds are grouped according to their classes indicated on the left. Higher relative resistance scores indicate higher resistance of PA14. All lactonases and AHLs are used at 50 μ M final concentrations, respectively.

### Screening of the data from the Biolog Phenotype MicroArray experiments

Results from the MicroArrays (**Fig. S1-S5**) were filtered, and this study focuses on compounds meeting all the following three criteria – (i) A differential change in PA14 growth is observed between the lactonase (SsoPox W263I and GcL) and AHL (C4-HSL and 3oC12-HSL) treatments at a given concentration. (ii) At this concentration, the antibiotic or antibacterial compound alters the growth of PA14 by at least 25% in at least one of the treatments (lactonase or AHL) compared to the control (5A8). (iii) The relative resistance score of PA14 against that antibiotic or antibacterial compound at that concentration should be 70% or lesser. These three filtering criteria eliminate compounds for which weak changes are observed and allow for the identification of molecules for which the resistance of PA14 is more likely to be regulated by QS signaling (**Fig. 1 and 2**). The first filter is self-explanatory, *i.e.,* different outputs for QS and QQ treatment may mean that a biological pathway or mechanism regulated by QS is involved in the observed change in resistance. The second filter stems from the observation that changes below 25% did not robustly replicate in new culture experiments described in the following section. The third criterion is based on the fact that changes in resistance are more easily and robustly observed when the antibiotics treatment substantially reduces growth (**Fig. S1-S5**).

### PA14 growth experiments with antibiotics

We replicated key observations from the phenotype MicroArray studies. For this purpose, we designed PA14 growth experiments using similar growth conditions on 96-well microplates. Growth, harvesting, and washing of PA14 were done similarly to described above. An inoculum solution is made by diluting the washed PA14 cells 1:100 into fresh Biolog IF-10A media. Sublethal concentrations of all the following tested antibiotics and antibacterial compounds were determined by performing dose-response experiments using a similar growth protocol as described above (**Fig. S8**). The compounds that we tested included – nafcillin (200 μg/mL), D-cyclo-serine (200 μg/mL), sulfathiazole (25 μg/mL), sulfadiazine (25 g/mL), μg/mL), trimethoprim (20 g/mL), procaine (500 μg/mL), carbonyl cyanide 3-chlorophenylhydrazone or CCCP (50 μg/mL) μg/mL). The growth of PA14 against these compounds at indicated sublethal and colistin (0.1 μg/mL), inactive lactonase dosages in the presence of lactonases (SsoPox W263I or GcL, 50 μg/mL), or pure exogenously added AHLs (C4-HSL or μM) was determined similarly to the Phenotype MicroArray experiments but conducted in replicates. 150 μL of each set of supplemented inoculum solution was dispensed into at least 4 consecutive wells of a 96 well sterile non-binding polypropylene microplate (Corning #3879) and topped with 50 μL sterile light mineral oil to prevent evaporation. The microplate was incubated and the growth of PA14 was recorded in a BioTek Epoch2 microplate spectrophotometer as described above.

### Total RNA extraction

Overnight cultures of PA14 in LB media were diluted 1:100 into fresh LB and grown until OD600 is ∼ 0.3 - 0.4. The cultures were harvested by centrifugation (5,000g for 3 min), and washed thrice with Biolog IF-10A media (without dye mix A). Washed cells were re- suspended in Biolog IF-10A media (without dye mix A) at an OD_750_ of ∼0.05. The washed cells were diluted 1:100 into 2 mL fresh Biolog IF-10A media (without dye mix A) with or without antibiotics sulfathiazole (25 μ (Corning #352059) and supplemented with 50 μ g/mL) in polypropylene culture tubes M (final) *N*-acyl homoserine lactones. Each condition was replicated 4 times. The cultures were incubated at 37°C for 22 h and shaken at 250 rpm. After incubation, the cultures were cooled on ice and the cells from 1 mL of culture were harvested by centrifugation (6000g for 5 min at 4°C). Cell pellets were frozen at -80°C until RNA extraction. Total RNA was extracted using the RNeasy mini kit (Qiagen #74104) following the manufacturer’s instructions. During purification, residual genomic DNA contamination was removed by an on-column DNAse digestion performed using the RNAse-free DNAse I kit (Qiagen #79254) following the manufacturer’s protocol at 37°C for 1 hr. The quality and quantity of purified RNA were tested using the Take3 plates on a BioTek Synergy HTX microplate reader. All extracted RNA was stored at -80°C until use.

### Quantitative Reverse Transcription Polymerase Chain Reaction (qRT-PCR)

qRT-PCR was performed using the Power SYBR® Green RNA-to-CT™ 1-Step Kit (Thermo Fisher Scientific #4389986) on a StepOnePlus™ Real-Time PCR System (Thermo Fisher Scientific) following the manufacturer’s instructions. RT-PCR grade water (Ambion #AM9935) was used for setting up all reactions. Primers for PA14 *folA*, *folP,* and *recA* genes (**Table S1**) were initially characterized to determine optimal concentrations and efficiencies in qPCR reactions using the Thermo Fisher Scientific StepOne^TM^ software v2.3. Total reaction volumes were 10 concentrations used were – *folA* (100 μL. Primer M forward and reverse primers), and *recA* (100 μM forward and reverse primers). Appropriate qRT-PCR controls – no template, no primer, and no reverse-transcriptase controls were performed for each reaction set. A relative quantification approach was undertaken to determine the amounts of *folA* and *folP* mRNA in each reaction, by using *recA* as the endogenous control gene.

### Caenorhabditis elegans liquid killing assays

Some experimental conditions changes were required to adapt the PA14 antibiotic resistance study conditions to the *C. elegans* infection model, including the growth media and incubation temperature. Standardized liquid killing assay [67] requires a specific culture medium to co-incubate the nematodes with PA14 at a temperature not exceeding 25°C, as that would otherwise be lethal for the nematodes. We noted that the Biolog IF-10A medium is toxic to *C. elegans* upon prolonged incubation, but that the toxicity can be reduced significantly with dilution (data not shown). Therefore, a 1:1 dilution of IF-10A media (without dye mix A) in the M9W buffer was chosen. It increases the mortality of *C. elegans* by 10-30% compared to a standard *C. elegans* maintenance buffer such as M9W in a 24-48 h incubation period (data not shown). We note that other buffer media, such as S-basal, S-complete, or SK media [68] could not be used because they caused the precipitation of the IF-10A media upon mixing.

### Growth of *C. elegans*

Strain SS104 [glp-4(bn2)] was obtained from the *Caenorhabditis* Genetics Center (CGC), University of Minnesota. The glp-4(bn2) mutation makes the nematodes incapable of producing offspring at temperatures greater than 20°C [69]. This is necessary to prevent the formation of progeny nematodes during the incubation period which will eventually hinder the downstream counting process. Nematodes were routinely maintained and cultured in the laboratory at 16°C on Nematode Growth Media (NGM) using standard protocols [68]. We modified a liquid killing assay protocol [67, 68] for this study. Synchronization of SS104 nematodes was performed by hypochlorite isolation of eggs from gravid adults. The eggs were washed and hatched in M9W medium for 24 h at 23°C to generate starved L1 larvae. Larvae were collected by centrifugation (1000g during 1min) and seeded on a 6 cm NGM-agar plate with a lawn of *E. coli* strain OP50 as food for the nematodes. The plate was incubated at 23°C for 48 h to generate a synchronized population of young adult nematodes. The nematodes were washed off the plate using M9W, followed by 6 subsequent wash steps interspersed with gravity-mediated settling of nematodes, and finally suspended in M9W. Work with *C. elegans* does not require IACUC approval.

### Growth of bacteria

Overnight cultures of *E. coli* OP50 and *P. aeruginosa* PA14 in LB media were diluted 1:100 into fresh LB and grown until OD_600_ is ∼ 0.3 - 0.4. The cultures were harvested by centrifugation (5000g for 3 min), washed thrice with M9W, and resuspended in M9W at a final OD_600_ of 0.3.

### Assay conditions

Young adult SS104 nematodes were co-incubated with *P. aeruginosa* PA14 in a 1:1 mix of IF-10A media (without dye mix A) and M9W buffer for 24 hrs. at 23°C with antibiotics (5 μ(100 μg/mL nafcillin or 25 μg/mL sulfathiazole) and lactonases μM). All components of the assay were mixed in a sterile flat-bottom L of assay medium, 96-well microplate (Sarstedt # 82.1581.001). Each well contained 200 μ composed of a mixture of M9W and Biolog IF-10A media (without dye mix A) at a ratio of 1:1, and supplemented with 10 μg/mL cholesterol, antibiotics, bacteria at a final OD_600_ of 0.03, *C. elegans* (at least 20 - 50 young adult nematodes per well). For experiments with PA14 only, enzymes (100 μM) were also added. As a control experiment, young adult nematodes were incubated with *E. coli* OP50 in the same media without antibiotics and lactonases/AHLs for a similar duration. OP50 is used as a standard food source for *C. elegans* during its routine laboratory maintenance and propagation and therefore, mortality observed when the nematodes are incubated with OP50 is treated as a control observation. For wells with OP50, sterile PTE buffer was added as a proxy for enzymes. At least 3 replicates were performed for each condition that was tested. Due to the inherent variability in manually pipetting nematode suspensions [70], the number of nematodes dispensed in each well varied, but each well contained at least 20, and most wells contained 20 to 50 nematodes. The microplate was then sealed with gas-permeable Breathe-Easy Membranes (Diversified BioTech #BEM-1) and incubated in a humidified incubator at 23°C for 24 h with shaking (300 rpm). After a 24 hr co-incubation period, the nematodes were allowed to settle at the bottom ofthe wells by gravity. 100 μ L containing the nematodes were transferred to unseeded 3.5 cm NGM-agar plates. The plates were sealed with parafilm and incubated at 23°C for a 24 hr recuperation period. The nematodes were then manually scored alive or dead under a microscope (Leica M165C) after touching them using a traditional worm pick [68].

### Data Analysis

Experimental data is represented as a % of nematodes that are alive after the entire incubation period for each treatment condition.

### Graphing and Analysis of Data

All data processing, analysis, and subsequent graphing is done either using Microsoft Excel or GraphPad Prism 8. Unpaired two-tailed t-tests with Welch’s correction were used for determining statistical significance and calculated using GraphPad Prism.

## Results and Discussion

The effect of QS signal disruption in *P. aeruginosa* was previously shown to be synergistic with some antibiotics, namely ciprofloxacin [29-31,35], ceftazidime [35] and gentamicin [29]. However, it remains unclear if this approach can be synergistic with all antibiotic therapies. To address this issue, we investigated if QQ can alter the global antibiotic resistance profile of *P. aeruginosa*. We used the Biolog Phenotype MicroArrays [66] to characterize the antibiotic-resistance profile of *P. aeruginosa* in the presence of QQ lactonases. While these MicroArrays have been previously used to study the metabolic characteristics of various *P. aeruginosa* strains under a range of conditions [66,71–76], or to study the chemical resistance in other bacteria [77–79] or mixed microbial communities [80], a comprehensive study of the antibiotic-resistance profile of laboratory strains of *P. aeruginosa* as a function of AHL signaling was not reported to our knowledge.

### Screening with antimicrobials reveals that resistance is modulated by lactonase treatment

We quantified the growth of PA14 using 10 Phenotype MicroArrays (Biolog PM11 to PM20) containing a total of 222 unique antibiotics and antibacterial compounds, in the presence of 4 experimental treatments – two lactonases (SsoPox W263I, GcL) and two pure exogenously added AHLs (C4-HSL, 3oC12-HSL) – and 1 control treatment, the inactive lactonase SsoPox 5A8 mutant (**Fig. 1-2; Fig. S1-S5**). An increase in the growth of PA14 in the presence of antimicrobial compounds relates to an increase in *resistance* (or a decrease in *sensitivity*) against these compounds. Conversely, decreased growth relates to an increase in *sensitivity* (decrease in *resistance*) against these compounds. It is in this context that the terms “sensitivity” and “resistance” are used throughout this manuscript.

Growth patterns of PA14 in the presence of antibiotics or antibacterial compounds appear considerably altered with lactonases/AHLs, and these alterations are also a function of the concentrations of these compounds (**Fig. S1-S5**). As expected for most of the antibiotics and antibacterial compounds, the growth of PA14 decreased with the increasing concentration of the compounds. The whole screening results shown in **Fig. S1-S5** were reduced using a set of criteria described in the Methods section to focus on the most robust changes upon treatments. Compounds of particular interest are shown in **Fig. 1 and 2**.

Results shown in **Figs. 1, 2, S1-S5** show that the antibiotic sensitivity profile of PA14 is dependent on QS signaling. Indeed, these tested antibiotics and antibacterial compounds can be broadly classified into two major groups. The first group (**Fig. 1**) includes conditions where lactonase treatment suppresses and/or exogenous AHLs promote the growth of PA14. QQ increases the sensitivity (or decreases the resistance) of PA14 against this group of molecules. The second group of tested antibiotics and biocides shows the opposite trend (**Fig. 2**): for these molecules, the lactonase treatments promote and/or exogenous AHLs suppress the growth of PA14, suggesting that QQ decreases the sensitivity (or increases the resistance) of PA14. As expected, changes in sensitivity of PA14 are more easily observed at antibiotic concentrations that exhibited a lower relative resistance score (**Fig. S1-S5**), usually below 70%.

Among the tested compounds that did not reduce PA14 growth much (resistance score >70%), a change in growth upon treatment (>25% in at least one treatment compared to 5A8 control) was observed for 10 molecules (**Fig. S6**). The sensitivity of PA14 to all these 10 compounds was significantly increased by QQ lactonase treatment, as compared to the control. For example, in the presence of chlorhexidine, PA14 showed a 19% and 42% reduction in growth upon QQ with SsoPox W263I and GcL, respectively, and a 9% and 11% increase in growth upon the addition of C4-HSL and 3oC12-HSL, respectively, compared to SsoPox 5A8 treated control. These types of observations are likely outliers and are largely eliminated using our filtering criteria (discussed in Methods).

We used the lactonases SsoPox W263I and GcL for AHL signal disruption. These enzymes were enzymatically and structurally characterized in previous studies [43-45,49,59]. Specifically, SsoPox shows a strong AHL preference for long acyl chain AHL substrates (>C8) and low activity against C4-HSL. On the other hand, GcL exhibits broad substrate preference and hydrolyzes both C4-HSL and 3oC12-HSL with high proficiency. The difference in kinetic properties of these two enzymes was previously described [59]. Since *P. aeruginosa* simultaneously utilizes two QS circuits based on C4-HSL and 3-oxo-C12 HSL, these two lactonases with distinct specificities may differentially affect these circuits. Differential QQ with these enzymes was previously described and resulted in differential proteome profiles, virulence factors expression, virulence, and biofilm formation in *P. aeruginosa* [56, 59].

To assess whether changes in antibiotic resistance would be sensitive to the differential quenching of AHL QS circuits, we focused on compounds for which robust resistance changes can be observed (see methods) and for which SsoPox W263I and GcL treatments show large differences (**Fig. S7**). For 27% of all tested compounds (60/222), changes in resistance upon lactonase treatment are unidirectional i.e., both lactonases either increase (32/60) or decrease (28/60) growth compared to control. Within these groups, some differences can be observed. For example, for a fraction of the compounds (5% of all tested compounds; 11/222), SsoPox W263I treatment resulted in >25 percentage points higher PA14 growth increase than GcL treatment (**Fig. S7**, top heat map panel). For another fraction (4.5% of all tested compounds; 10/222), GcL treatment yielded >25 percentage points higher PA14 growth compared to SsoPox W263I (**Fig. S7**, middle heat map panel). For another group of compounds (5.4% of all tested compounds; 12/222), treatment with both lactonases resulted in opposite changes in resistance, *i.e*. the change in the growth of PA14 was opposite with SsoPox W263I and GcL by >25% for at least one lactonase compared to 5A8 treated control (**Fig. S7**, bottom heat map panel). These observations suggest that the AHL-dependent changes in the resistance profile of PA14 may be sensitive to the AHL substrate specificity of the QQ lactonase.

### Replication experiments show that interference in AHL signaling induces changes in resistance that are variable in sign and magnitude

Observations derived from the experiments with the Biolog Phenotype MicroArrays were replicated for some key candidate compounds. First, dose-response experiments against PA14 were performed (**Fig. S8**) and used to determine their sublethal concentration values. This was needed because of the proprietary conditions of Biolog MicroArrays. The results (**Fig. 4**) confirm that the antibiotic resistance profile of PA14 is dependent on AHL signaling. Similar to the screening experiment, the same two groups of compounds can be established – (i) antibiotics for which lactonases suppress and/or exogenous AHLs promote the growth of PA14 (e.g., nafcillin, oxacillin, D-cyclo-serine, norfloxacin, and ofloxacin) and (ii) antibiotics for which lactonases promote and/or exogenous AHLs suppress the growth of PA14 (e.g., azlocillin, sulfadiazine sulfathiazole, and carbenicillin). Lactonase treatment increased the sensitivity of PA14 against nafcillin, oxacillin, D-cyclo-serine, and norfloxacin by up to 55%, 50%, 70% and 35% respectively, compared to control. On the contrary, lactonase treatment increased the resistance of PA14 against azlocillin, sulfadiazine, sulfathiazole, and carbenicillin by up to 176%, 73%, 215% and 362% respectively, compared to control. In the case of ofloxacin, no significant effect of lactonase treatment on PA14 sensitivity was observed. However, the addition of exogenous AHLs to stimulate QS increased the resistance of PA14 to ofloxacin by up to 23% compared to the control. These results suggest a complex relationship between QS and antibiotic resistance in PA14. For example, lactonase treatments and the addition of C4- HSL with antibiotics like sulfadiazine, sulfathiazole, and trimethoprim, increased the resistance of PA14 to these compounds by 25%, 15% and 28% respectively, compared to control (**Fig. 4**). This is not the case with 3oC12-HSL treatment, for which resistance of PA14 does not differ from control. Treatment with GcL increased the sensitivity and with C4-HSL increased the resistance of PA14 against trimethoprim by 21% and 28% respectively, compared to control (**Fig. 4**). This is consistent with the lactonase substrate specificity, because GcL degrades C4- HSL proficiently (and SsoPox W263I does not), therefore GcL and addition of C4-HSL appear to show antagonistic effects.

For some compounds, including procaine and coumarin, the results are surprising: both lactonase and AHL treatments increase the sensitivity of PA14 against these compounds by up to 51% (**Fig S11**). Coumarin was previously reported to be an inhibitor of QS circuits and additional signaling pathways in *P. aeruginosa* [81, 82], and this is expected to affect the results of our experiments. In the case of procaine, it was previously shown to enhance antibiotic resistance in *P. aeruginosa* by increasing the expression of the MexCD-OprJ and MexAB-OprM efflux pumps [83], and is therefore also likely to affect the outcome of our experiments.

Overall, most observations from these replicated experiments are consistent with results from the screening (**Fig. S9-S11**). However, some differences can be noted for a few compounds (e.g., ofloxacin, procaine, CCCP, colistin, and trimethoprim). These discrepancies likely originate from the sensitivity of QS-dependent regulations to growth conditions. The latter could not be exactly reproduced due to the proprietary composition of Biolog conditions. The fact that QS-dependent regulations are highly sensitive to growth conditions and treatment is illustrated by two different sublethal concentrations (150 and 200 μg/mL) of D-cyclo-serine (**Fig S12**). With 150 μg/mL of D-cyclo-serine, lactonase treatments increased PA14 sensitivity by 9%, while it increased PA14 sensitivity up to 70% with 200 μg/mL of D-cyclo-serine.

Additionally, we note that this increased sensitivity may originate from the positive regulation of alanine racemase in *P. aeruginosa* by QS, a target of D-cyclo-serine (see Table S1 in ref [84] for studies conducted with strain PAO1).

Altogether, these experiments (**Fig. 1, 2, S1-S5)** show that: (i) AHL mediated QS signaling can significantly alter the antibiotic resistance profile of *P. aeruginosa*; and (ii) changes in resistance are variable in sign and magnitude, in ways that are not inferable from the properties of the tested antibiotics and antibacterial compounds.

### The sign of the effect of lactonase treatment on resistance varies for the different tested antimicrobial groups

We classified the tested antibiotics and antibacterial compounds according to their type, class, or mode of action – namely aminoglycoside, antifolate, β lactam penicillin, quinolone and fluoroquinolone, glycopeptide, macrolide and lincosamide, nitrofuran, polymyxin, quaternary ammonium salt, rifamycin, sulfonamide, tetracycline, and inorganic salt. Antibacterial compounds not fitting into any of these categories were labeled as unclassified organic compounds. Because the bacterial resistance mechanisms developed are often conserved and specific to certain antibiotic or antibacterial compounds, we hypothesized that observed changes may be shared by compound classes. This is corroborated by previous observations showing that QQ enzymes can alter the levels of proteins typically involved in antibiotic resistance in PA14 proteomics studies [59].

Consistently, we observe groups of molecules for which QQ lactonase treatment is either neutral or increases the sensitivity of PA14 (**Fig. 3**; i.e., antifolates, polymyxins, rifamycins, quinolones, fluoroquinolones, nitrofurans, macrolides, lincosamides, and quaternary ammonium compounds). We also observe groups of molecules for which QQ lactonase treatment is either neutral or decreases the sensitivity of PA14 (**Fig. 3**; sulfonamides, tetracyclines). Other groups, for which QS modulation can either increase or decrease resistance, include inorganic salts, an unclassified group of organic compounds, and the –β lactam group (penicillins and cephalosporins). We note that for most conditions, QS alteration resulted in no change in PA14 resistance. This may be caused by the fact that the involved pathways are unaffected by quorum quenching, and/or that the inherent resistance of PA14 against these compounds and the fact that tested concentrations were too low to sufficiently challenge PA14 growth. This is illustrated by the observed correlation between the relative resistance score and the magnitude of the observed modulations by QS alterations.

**Figure 3.**
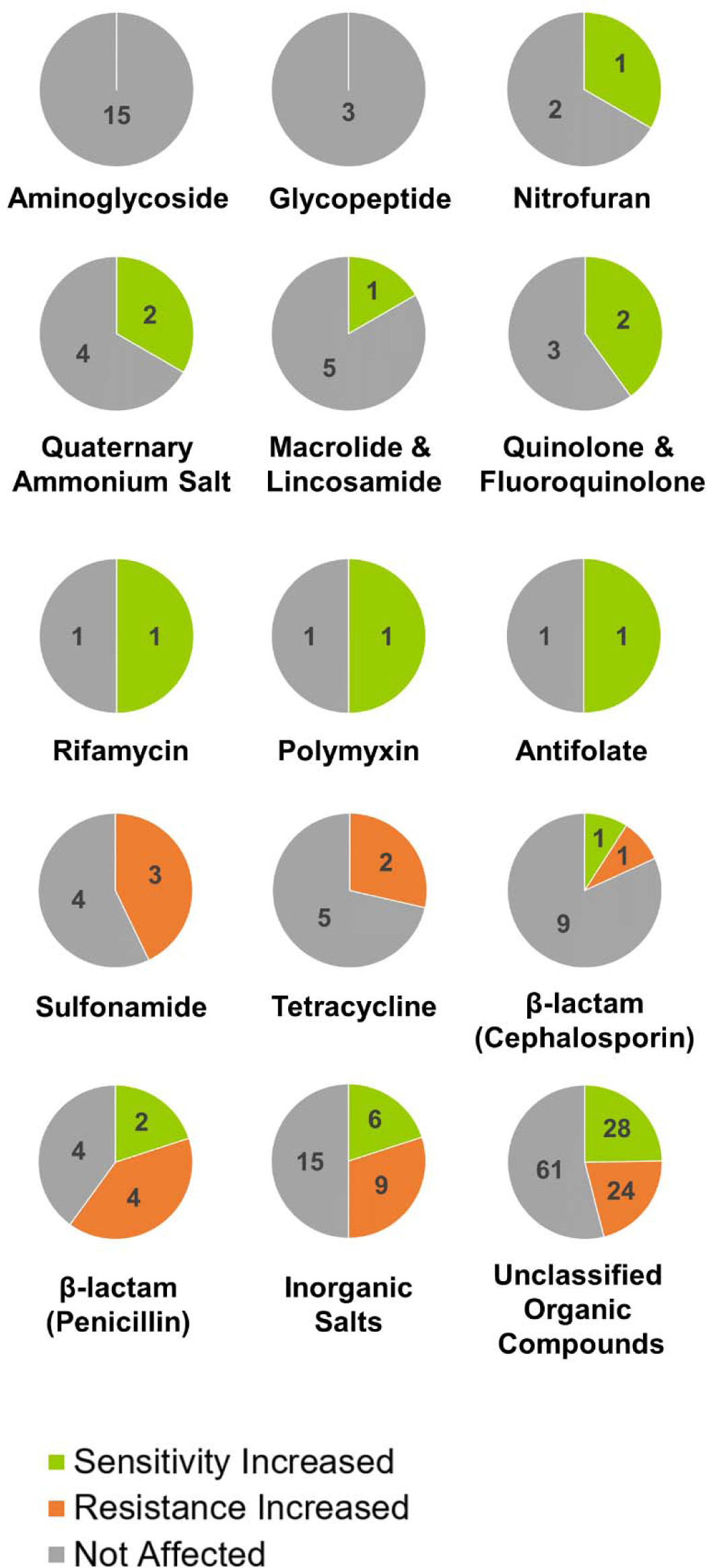
Effects of AHL signal disruption on different classes of antimicrobials. Pie charts showing the fraction of tested compounds from different classes for which a change in sensitivity and/or resistance of PA14 upon AHL signal disruption is observed.

AHL signal disruption by lactonases led to exclusively increased sensitivity of PA14 to 40% of tested fluoroquinolones (2/5) on the Biolog Phenotype MicroArrays (**Fig. 3**). Fluoroquinolone antibiotics inhibit bacterial DNA synthesis by targeting DNA topoisomerases essential for DNA replication in bacteria – DNA gyrase (in Gram-negative bacteria) and topoisomerase IV (in Gram-positive bacteria) [85]. Typical mechanisms of resistance against this class of antibiotics involve antagonistic mutations in target topoisomerases, use of efflux pumps to secrete the drugs out of the cell, or reducing their membrane permeability through porins [86]. In the case of *P. aeruginosa*, the RND efflux pumps MexAB-OprM [87], MexCD-OprJ [88], and MexEF-OprN [89] were implicated in fluoroquinolone resistance. Here, the increased sensitivity of PA14 observed with lactonase treatment might be related to decreased OprM levels upon lactonase treatment as was previously reported [59]. Conversely, adding exogenous AHLs increases resistance to ofloxacin (**Fig. 4**). This is consistent with a previous report of a lasR overproducing *P. aeruginosa* strain found to be more resistant to ofloxacin, and in the absence of lasR (QS compromised), the overexpression of master regulator RpoS can restore the loss of ofloxacin resistance [90]. We also observe that lactonase treatment with GcL (but not SsoPox) increased its sensitivity towards norfloxacin (**Fig. 4**). This resonates with other studies reporting that the loss of function of the regulator protein NfxB, which represses the production of MexCD-OprJ efflux pump, is associated with increased resistance of *P. aeruginosa* against norfloxacin [91, 92] and is under the regulatory control of the master regulator VqsM that promotes QS by producing LasI an (enzyme that synthesizes 3-oxo-C12- HSL in PA14) [93]. Additionally, lactonase-mediated QQ also had a synergistic effect when co-administered with the fluoroquinolone antibiotic ciprofloxacin and prevented the spread of *P. aeruginosa* in a burn wound infection model [30].

**Figure 4.**
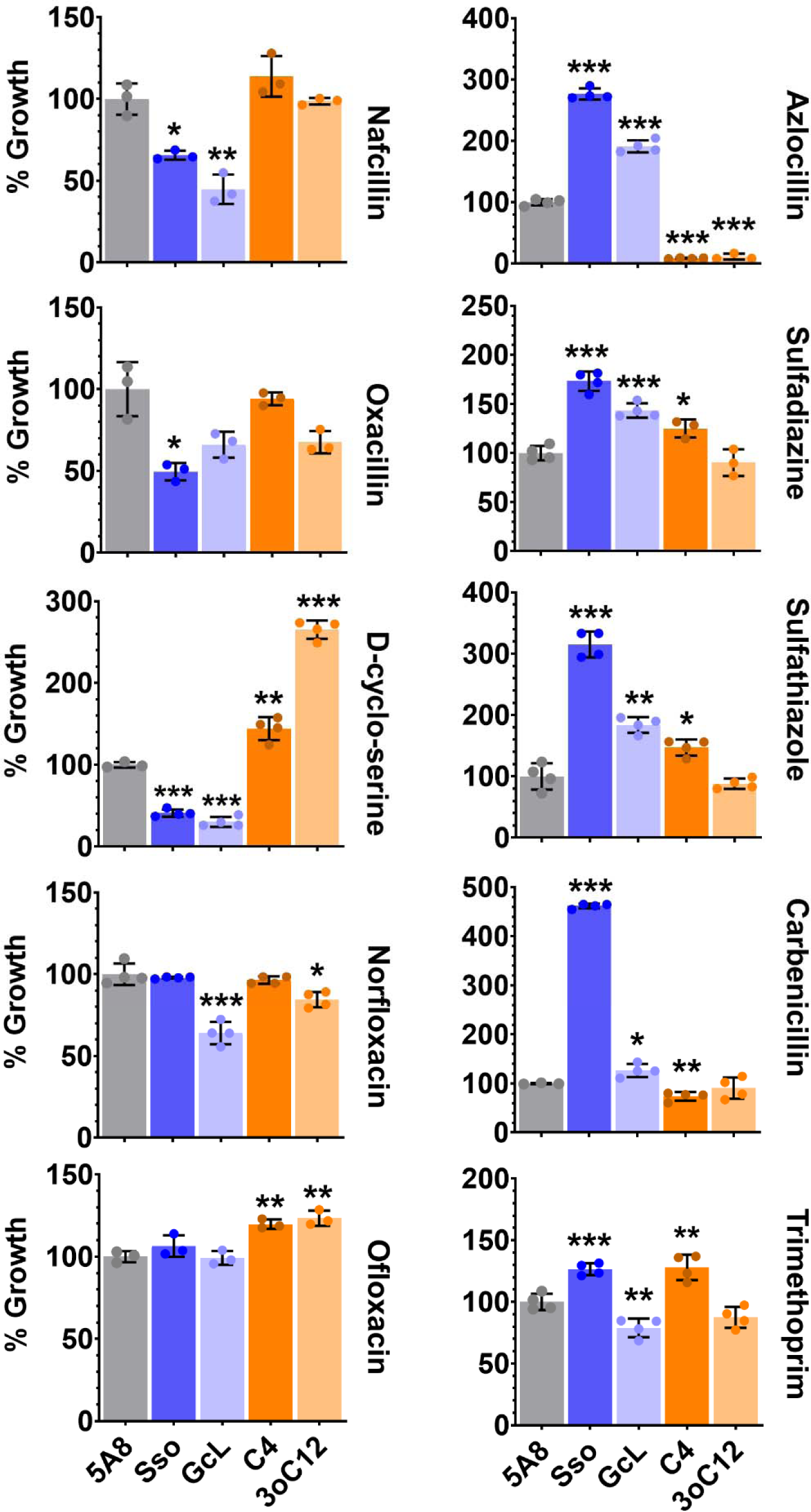
Replicated PA14 growth experiments in the presence of 10 antimicrobials identified in the screening experiment and interference in quorum sensing compounds. Treatments are: QQ lactonases – SsoPox W263I (Sso) and GcL or exogenously added AHLs – C4-HSL (C4) and 3-oxo-C12-HSL (3oC12), compared to control treatment (inactive lactonase SsoPox 5A8 (5A8)). PA14 growth in all treatments is normalized to the respective 5A8 control. All lactonases and AHLs are used at 50 μ Following concentration of antibiotics were used – 200 μM final concentrations, respectively. g/mL oxacillin, 200 g/mL azlocillin, 25 g/mL μ g/mL trimethoprim. All sulfadiazine, 25 μ g/mL carbenicillin and 20 μexperiments were done, and all data is represented as the mean and standard deviation of at least triplicates. Statistical significance of all treatments compared to the control (5A8) was calculated using unpaired two-tailed t-tests with Welch’s correction and significance values are indicated as - \*\*\**p < 0.0005*, \*\**p < 0.005* and \**p < 0.05*.

For other groups, such as β-lactams, alteration of QS modulation can either increase or β decrease resistance. When AHL signaling is inhibited by lactonases, *P. aeruginosa* becomes more sensitive to nafcillin and oxacillin but more resistant to azlocillin and carbenicillin (**Fig. 4**).

This discrepancy is possibly a result of the use of different resistance mechanisms against these drugs. While *P. aeruginosa* strains were reported to produce β-lactamases such as AmpCβ [94] and OXA-50 [20] that offer protection against a broad range of β-lactam antibiotics, the β resistance against some classes of β-lactam antibiotics can be mediated by the regulation of the β expression of outer membrane porins and efflux pumps such as OprD and MexXY-OprM [95]. In support of this hypothesis, a previous proteomics study reported an increase in OprD and a decrease in OprM levels in PA14 upon AHL signal disruption using lactonases SsoPox W263I and GcL [59]. A synergy between nafcillin and QQ agents was also previously reported for Gram-positive pathogens – *Staphylococcus aureus* [96] and *Staphylococcus epidermidis* [97].

### AHL-mediated quorum sensing and resistance of *P. aeruginosa* to sulfathiazole and trimethoprim

Sulfathiazole and trimethoprim target the enzymes involved in the biosynthesis of folates in bacteria [98]. Sulfathiazole belongs to a class of compounds called sulfonamides that act as competitive inhibitors of the enzyme dihydropteroate synthase (DHPS; encoded by the *folP* gene in PA14). Trimethoprim, an antifolate compound, targets the enzyme dihydrofolate reductase (DHFR; encoded by the *folA* gene in PA14) [99]. While DHPS acts upstream of the folate biosynthesis pathway converting *p*-amino benzoic acid to dihydropteroate, DHFR is the enzyme catalyzing the terminal step of the pathway leading to the reduction of dihydrofolate to tetrahydrofolate [100]. Bacteria can develop resistance against sulfonamides or trimethoprim by using several mechanisms [101, 102] - accumulation of compensatory mutations in the native DHPS/DHFR enzymes that prevent binding of the drugs [103, 104], using drug efflux pumps and altering membrane barrier permeability [105, 106], horizontal acquisition of foreign drug-resistant DHPS/DHFR variants or homologs either on the chromosome via mobile genetic elements or via plasmids [107, 108], genetic regulation of DHPS/DHFR production [109, 110], or producing specialized enzymes that cleave these drugs [111].

In this study, the modulations of the resistance to sulfonamide and trimethoprim observed in *P. aeruginosa* PA14 by QS are likely to originate from the genetic regulation of *folA* and *folP* genes. To test this hypothesis, we carried out qRT-PCR experiments to determine the expression levels of *folA* and *folP* transcript mRNA under a variety of conditions (**Fig. 5**). In the absence of any antibiotics, when QS is attenuated by lactonases, *folA* (**Fig. 5A**) and *folP* (**Fig. 5B**) genes are upregulated by 4-fold and 3-fold, respectively, compared to an inactive enzyme treated control. This suggests a possible increased metabolic demand for folate when QS is suppressed. When AHL signaling is attenuated and PA14 is challenged with sulfathiazole, expression of *folA* remains significantly upregulated (between 2 to 2.5-fold; **Fig. 5C**) compared to the inactive enzyme treated control. However, *folP* expression is downregulated by 30% (**Fig. 5D**). In the presence of trimethoprim, only *folP* expression is increased (2-fold, **Fig. 5F**) as compared to control. This downregulation of *folP* upon AHL signal disruption is also observed in the absolute levels of transcripts (**Fig. S13**). Under most conditions (**Fig. S13**), as expected, competitive inhibitors like sulfathiazole and trimethoprim increase the production of FolA and FolP enzymes, compared to the no-antibiotic control. However, when AHL signaling is disrupted by lactonases, the absolute *folP* levels in sulfathiazole treated PA14 are reduced to similar levels to those observed for untreated PA14.

**Figure 5.**
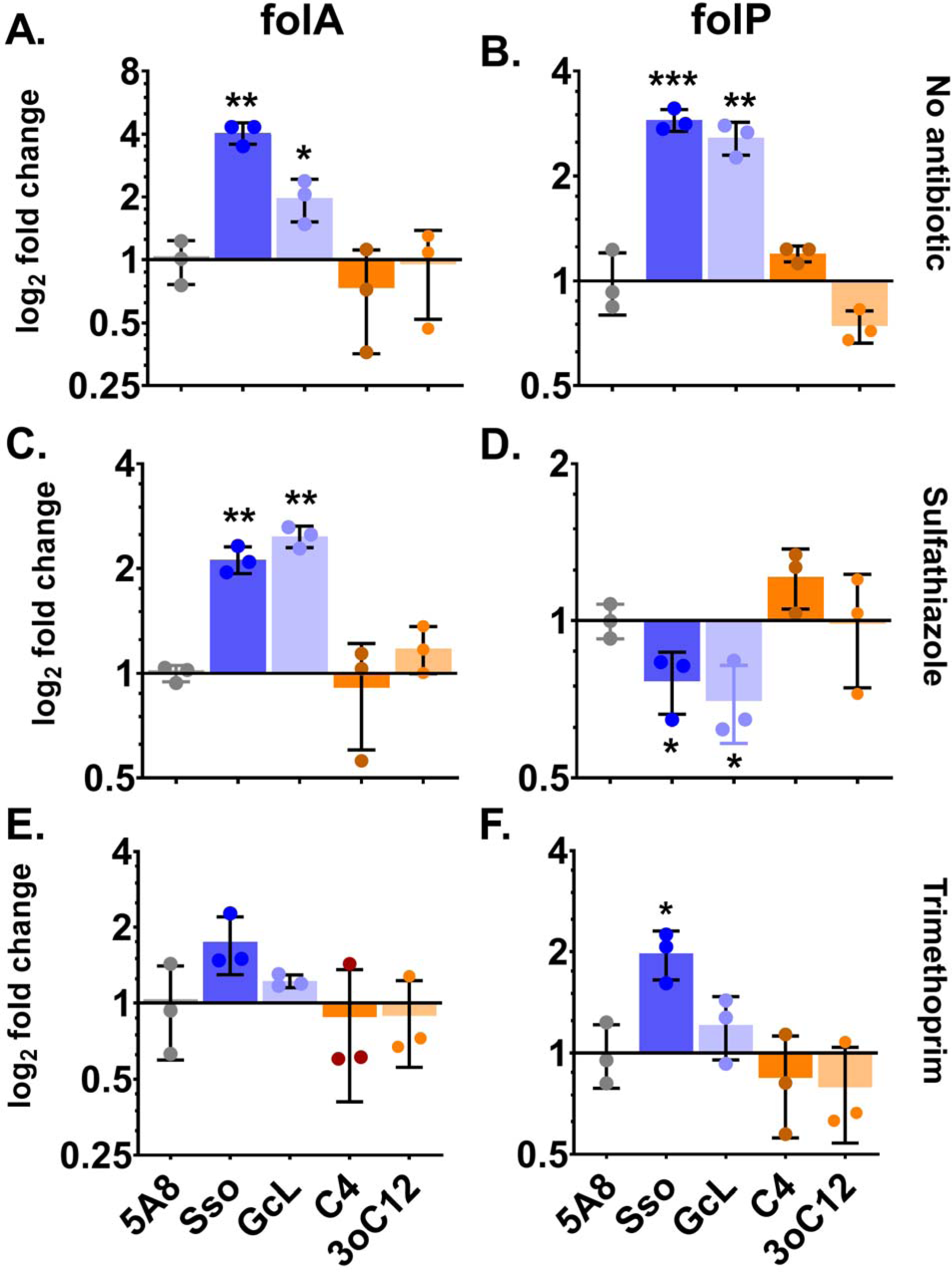
Changes in mRNA levels (as measured by qRT-PCR) of genes folA and folP with sulfathiazole and trimethoprim and as a function of quorum sensing interference. Used treatments are no antibiotics (A, B) or with antibiotics sulfathiazole (C, D) and trimethoprim (E, F). Tested conditions include the QQ lactonases – SsoPox W263I (Sso) and GcL or exogenously added AHLs – C4-HSL (C4) and 3-oxo-C12-HSL (3oC12), compared to control treatment (inactive lactonase SsoPox 5A8 (5A8)). *folA* and *folP* mRNA levels in all treatments were determined using the relative quantification method using the *recA* gene as endogenous control and normalized to the respective 5A8 control, which is set to 1 on a log_2_ scale. All lactonases and AHLs were used at 50 μM final concentrations, respectively. The following concentration of antibiotics was used – 25 μg/mL trimethoprim. All experiments were done, and all data is represented as the mean and standard deviation of at least triplicates. Statistical significance of all treatments compared to the control (5A8) was calculated using unpaired two-tailed t-tests with Welch’s correction and significance values are indicated as - \*\*\**p < 0.0005*, \*\**p < 0.005* and \**p < 0.05*.

These observations suggest that QS regulates the expression of *folP* and *folA* genes in PA14. We also observe that QQ leads to increased resistance of PA14 to sulfathiazole (**Fig. 5**). In a study previously conducted in yeast auxotrophs [109], sulfonamides, in addition to inhibiting DHPS, are metabolized to sulfa-dihydropteroate, a compound that inhibits the downstream enzyme DHFR and thereby the biosynthesis of folate. This inhibition can be thwarted by overproducing DHFR. When challenged with sulfathiazole and AHL signal disruption, we observed a simultaneous downregulation of *folP* (DHPS; **Fig. 5D**) and overproduction of *folA* (DHFR; **Fig. 5C**). This *folP* and *folA* regulation might explain the higher resistance of PA14 to sulfathiazole when QS is disrupted.

Unlike sulfathiazole, changes in resistance of PA14 against trimethoprim upon AHL signal disruption cannot be explained solely by the observed changes in *folP* and *folA* regulation. An increase in absolute (**Fig. S13**) and relative levels (compared to inactive lactonase 5A8 treatment, **Fig. 5F**) of *folP* in the presence of trimethoprim and lactonase can be observed, and it might contribute to the observed increased resistance (**Fig. 4**). Adaptive resistance mechanisms such as the regulation of efflux pumps might be involved in the QS-dependent regulation of trimethoprim resistance in PA14. For example, MexEF-OprN efflux pump overexpression (due to loss of function of its repressor NfxC) renders *P. aeruginosa* resistant to several antibiotics including trimethoprim [112]. Similarly, the BpeEF-OprC efflux pump system confers trimethoprim resistance to *Burkholderia pseudomallei* [113].

AHL signaling can modulate the potency of antibiotic treatments in a *P. aeruginosa C. elegans* infection model.

Upon observing that interference in AHL signaling modulates antibiotic resistance of *P. aeruginosa* in liquid cultures, we examined whether these modulations would translate into altered PA14 pathogenicity in an *in vivo Caenorhabditis elegans* infection model. *C. elegans* is a nematode whose genotypes and phenotypes have been characterized and have long served as a model organism in cellular, molecular, and developmental biology research [114, 115]. It has also been extensively used as a convenient model animal host in a plethora of microbial infection assays [68,116–118]. We adapted the standard liquid killing assay (see methods section). Lactonases were previously shown to reduce the virulence of *P. aeruginosa* against *C. elegans* in infection assays [46].

When challenged with nafcillin or sulfathiazole, the pathogenicity of PA14 in a *C. elegans* killing assay is significantly modulated by AHL-based quorum sensing (**Fig. 6A, 6B**). For instance, in the presence of nafcillin (**Fig. 6A**) and sulfathiazole (**Fig. 6B**), QQ lactonase treatment reduced the virulence of PA14 against *C. elegans*. Nematode survival increased by up to 34% and 150% for nafcillin and sulfathiazole respectively, upon treatment with SsoPox W263I and GcL compared to the inactive enzyme control. Conversely, with azlocillin, nematode mortality is increased by lactonase treatment by up to 81% compared to control (**Fig. 6C**).

**Figure 6.**
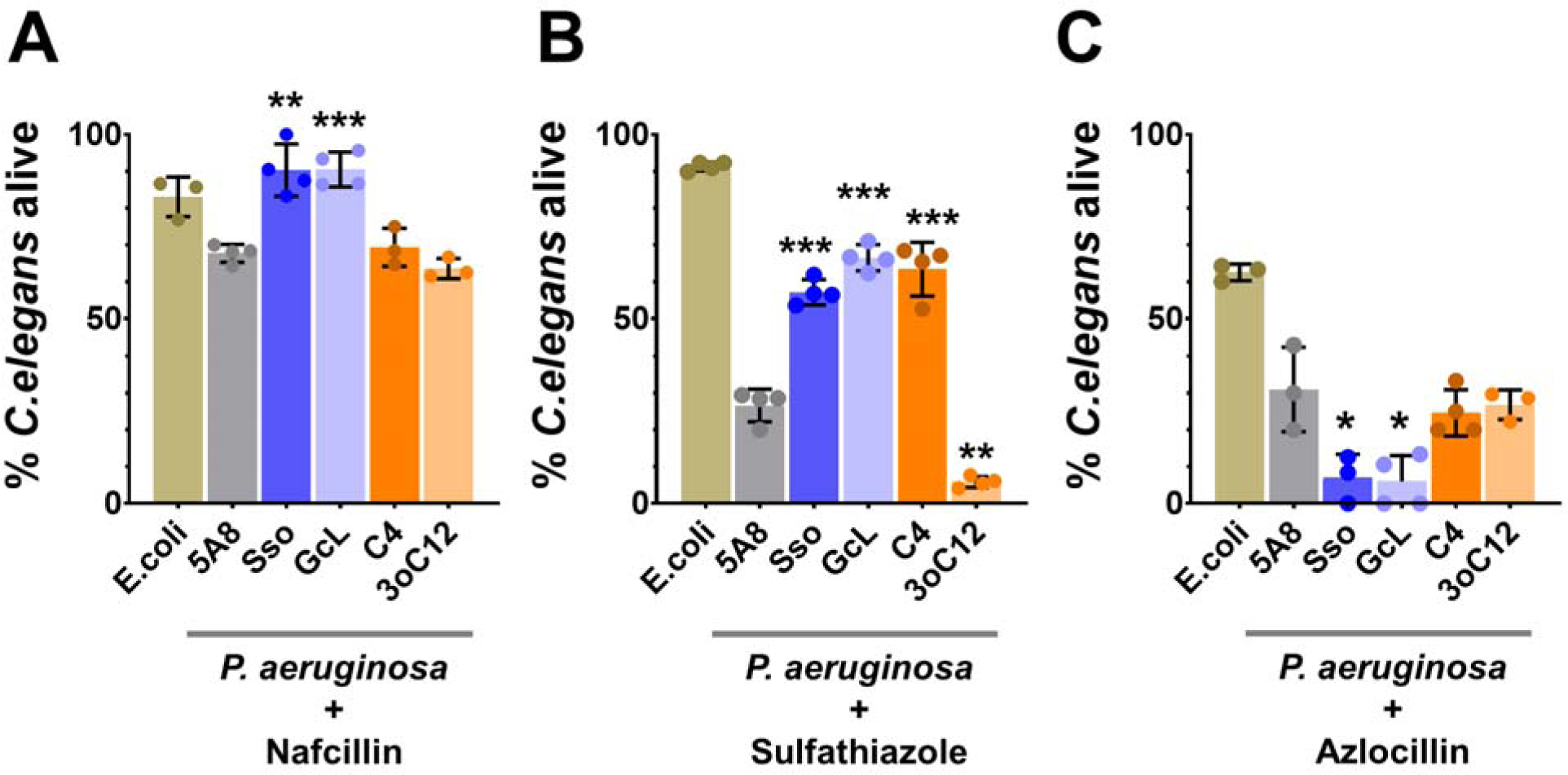
*In vivo C. elegans* infection to evaluate the effects of the interference in quorum sensing and antibiotic treatments. Assays were performed with antibiotics – nafcillin (A), sulfathiazole (B) and azlocillin (C) upon treatment with QQ lactonases – SsoPox W263I (Sso) and GcL or exogenously added AHLs – C4-HSL (C4) and 3-oxo-C12-HSL (3oC12), compared to control treatment (inactive lactonase SsoPox 5A8 (5A8)). Virulence of PA14 is exemplified by the death of *C. elegans* upon infection. Mortality is represented as the % of nematodes that survived the assay. *E. coli* strain OP50 is used as a non-virulent control for the assays. All lactonases and AHLs are used at 100 μconcentration of antibiotics were – 200 μg/Ml final concentrations, respectively. Used sulfathiazole and 5μg/Ml Lazlocillin. All experiments were done, and all data is represented as the mean and standard deviation of at least triplicates. Statistical significance of all treatments compared to the control (5A8) was calculated using unpaired two-tailed t-tests with Welch’s correction and significance values are indicated as - \*\*\**p < 0.0005*, \*\**p < 0.005* and \**p < 0.05*.

Alteration of the antibiotic resistance profile of PA14 *in vivo* by AHL-based QS is further evidenced by the observed modulations of the nematode mortality upon treatment with AHL signaling molecules. For example, treatment of PA14 with sulfathiazole and 3oC12-HSL caused an increase in nematode mortality (+78%), whereas sulfathiazole and C4-HSL treatment led to an increase in survival (+138%), compared to an inactive lactonase treated control. This may suggest that the regulation of virulence, in the presence of sulfathiazole, is similar to that described for pyocyanin production. Indeed, Rhl QS circuit agonists like C4-HSL were reported to suppress pyocyanin production in *P. aeruginosa* in a PQS-dependent manner [119, 120].

The changes in pathogenicity of PA14 against *C. elegans* upon treatment with nafcillin or azlocillin and QQ lactonases echo a trend observed in the PA14’s resistance profile against these compounds. QQ lactonase treatment increased the sensitivity of PA14 to nafcillin (**Fig. 4**), and its virulence against *C. elegans* is also reduced in the presence of nafcillin (**Fig. 6A**). Similarly, the resistance of PA14 to azlocillin is increased upon treatment with QQ lactonases (**Fig. 4**), and accordingly, its virulence against *C. elegans* is increased in the presence of QQ lactonases and azlocillin (**Fig. 6C**). Unexpectedly, however, we observe that while the treatment with QQ lactonases increases the resistance of PA14 to sulfathiazole (**Fig. 4**), its virulence is reduced (**Fig. 6B**), suggesting that the relationship between antibiotic resistance and its ability to kill the nematodes can be complex.

Taken together, these observations show that AHL-based QS and signal disruption can alter the pathogenicity of *P. aeruginosa* against *C. elegans* in the presence of antibiotics. QQ lactonases can modulate the sensitivity of PA14 to an antibiotic in a manner that may not always be reflected in changes in its virulence under similar conditions. Therefore, it appears that a combined administration of antibiotics and QQ agents can result in virulence changes that are difficult to predict and can either prove to be beneficial or detrimental to the host.

## Conclusion

In this study, we demonstrate that the disruption of AHL-mediated QS signaling by the thermostable lactonases SsoPox and GcL can alter the antibiotic resistance profile of the Gram-negative pathogen *P. aeruginosa* PA14. This likely occurs *via* changes in the regulation of both intrinsic and adaptive resistance mechanisms. We were able to translate the observations from the Biolog Phenotype MicroArrays into experimental replicates using similar growth conditions for multiple antibiotics at their respective sublethal concentrations. Changes in antibiotic resistance against sulfathiazole and trimethoprim could be linked to key changes in the expression levels of genes involved in folate biosynthesis. Lastly, these observations were evaluated with an *in vivo C. elegans* infection model, and the results confirm that (i) the ability of *P. aeruginosa* to kill the nematode in presence of certain antibiotics depends on QS; (ii) yet that the combined effects of antibiotics and QQ lactonase are not always synergistic. Overall, whereas most previous studies investigating the effect of co-administration of QQ agents (QS inhibitors, enzymes) and antibiotics in bacterial infection models reported a positive synergistic effect [30,37,97], our results suggest that in *P. aeruginosa*, the relationship between antibiotics and QQ agents is very complex, and depends on the type of antibiotic and substrate preference of the QQ agent used. More studies are needed to decipher the mechanisms for the synergies and antagonisms described in this work to assess and evaluate the potential of combination therapy in *P. aeruginosa*.

## Acknowledgement.

We would like to thank Dr. Eliana Drenkard and Dr. Frederick Ausubel at the Massachusetts General Hospital for providing us with *Pseudomonas aeruginosa* strain PA14, Dr. Barry Bochner at Biolog Inc. for helping us setting up the experimental protocol with Biolog Phenotype MicroArrays and Aric Daul at the *Caenorhabditis* Genetics Center (CGC) at the University of Minnesota for providing us with *Caenorhabditis elegans* strain SS104 and *Escherichia coli* strain OP50. This work was supported by the National Institute of General Medical Sciences of the National Institutes of Health under award number R35GM133487. The content is solely the responsibility of the authors and does not necessarily represent the official views of the National Institutes of Health.

