## Supplementary material for "Evidence for complex interplay between quorum sensing and antibiotic resistance in *Pseudomonas aeruginosa*"

---

### Table of Contents

**Page 2 and 3:** Table of contents.

**Page 4:** Table S-1: qPCR primers used in this study.

**Page 5:** Fig. S1A: Growth profile of *P. aeruginosa* PA14 with antimicrobials and interference in quorum sensing.

**Page 6:** Fig. S1B: Growth profile of *P. aeruginosa* PA14 with antimicrobials and interference in quorum sensing.

**Page 7:** Fig. S2A: Growth profile of *P. aeruginosa* PA14 with antimicrobials and interference in quorum sensing.

**Page 8:** Fig. S2B: Growth profile of *P. aeruginosa* PA14 with antimicrobials and interference in quorum sensing.

**Page 9:** Fig. S3A: Growth profile of *P. aeruginosa* PA14 with antimicrobials and interference in quorum sensing.

**Page 10:** Fig. S3B: Growth profile of *P. aeruginosa* PA14 with antimicrobials and interference in quorum sensing.

**Page 11:** Fig. S4A: Growth profile of *P. aeruginosa* PA14 with antimicrobials and interference in quorum sensing.

**Page 12:** Fig. S4B: Growth profile of *P. aeruginosa* PA14 with antimicrobials and interference in quorum sensing.

**Page 13:** Fig. S5A: Growth profile of *P. aeruginosa* PA14 with antimicrobials and interference in quorum sensing.

**Page 14:** Fig. S5B: Growth profile of *P. aeruginosa* PA14 with antimicrobials and interference in quorum sensing.

**Page 15:** Fig. S6: Heat maps representing growth of *P. aeruginosa* PA14 in presence of antibacterial compounds for which the sensitivity of PA14 is regulated by AHL signaling (see main text).

**Page 16:** Fig. S7: Modulations of the antibiotic resistance profile of *P. aeruginosa* PA14 depends on the AHL substrate specificity of QQ lactonase.

**Page 17:** Fig. S8: Dose-response growth of *P. aeruginosa* PA14 to determine the sublethal testing concentration for several antibiotics and antibacterial compounds.

**Page 18:** Fig. S9: Comparison between *P. aeruginosa* PA14 growth in Phenotype MicroArray (PM) cultures and replicated experiments with nafcillin, oxacillin, D-cyclo-serine, norfloxacin and ofloxacin.

**Page 19:** Fig. S10: Comparison between PA14 growth in Phenotype MicroArray (PM) cultures and replicated experiments with azlocillin, sulfadiazine, sulfathiazole, carbenicillin and trimethoprim.

**Page 20:** Fig. S11: Comparison between *P. aeruginosa* PA14 growth in Phenotype MicroArray (PM) cultures and replicated experiments with procaine, coumarin, CCCP and colistin.

**Page 21:** Fig. S12: *P. aeruginosa* PA14 growth in replicated experiments with antibiotics are sensitive to changes in the sublethal concentration of the antibiotic.

**Page 22:** Fig. S13: qRT-PCR quantifications of changes in transcript mRNA levels of genes folA and folP.

**Table S1: qPCR primers used in this study.**

| PA14 Gene | Identifier | Sense | Sequence |
|---|---|---|---|
| folA | PA14_04580 | forward | 5'-GATGATCGCCGCCCTTG |
| folA | PA14_04580 | reverse | 5'-GAGGGTCATCGCCTTGAAAT |
| folP | PA14_62850 | forward | 5'-GCATGATCGGCAAGGTACT |
| folP | PA14_62850 | reverse | 5'-AATTATCCGTGCGCCCTT |
| recA | PA14_17530 | forward | 5'-CGCAAGATCACCGGCAATATCA |
| recA | PA14_17530 | reverse | 5'-GGACCGAGGCGTAGAACTTC |

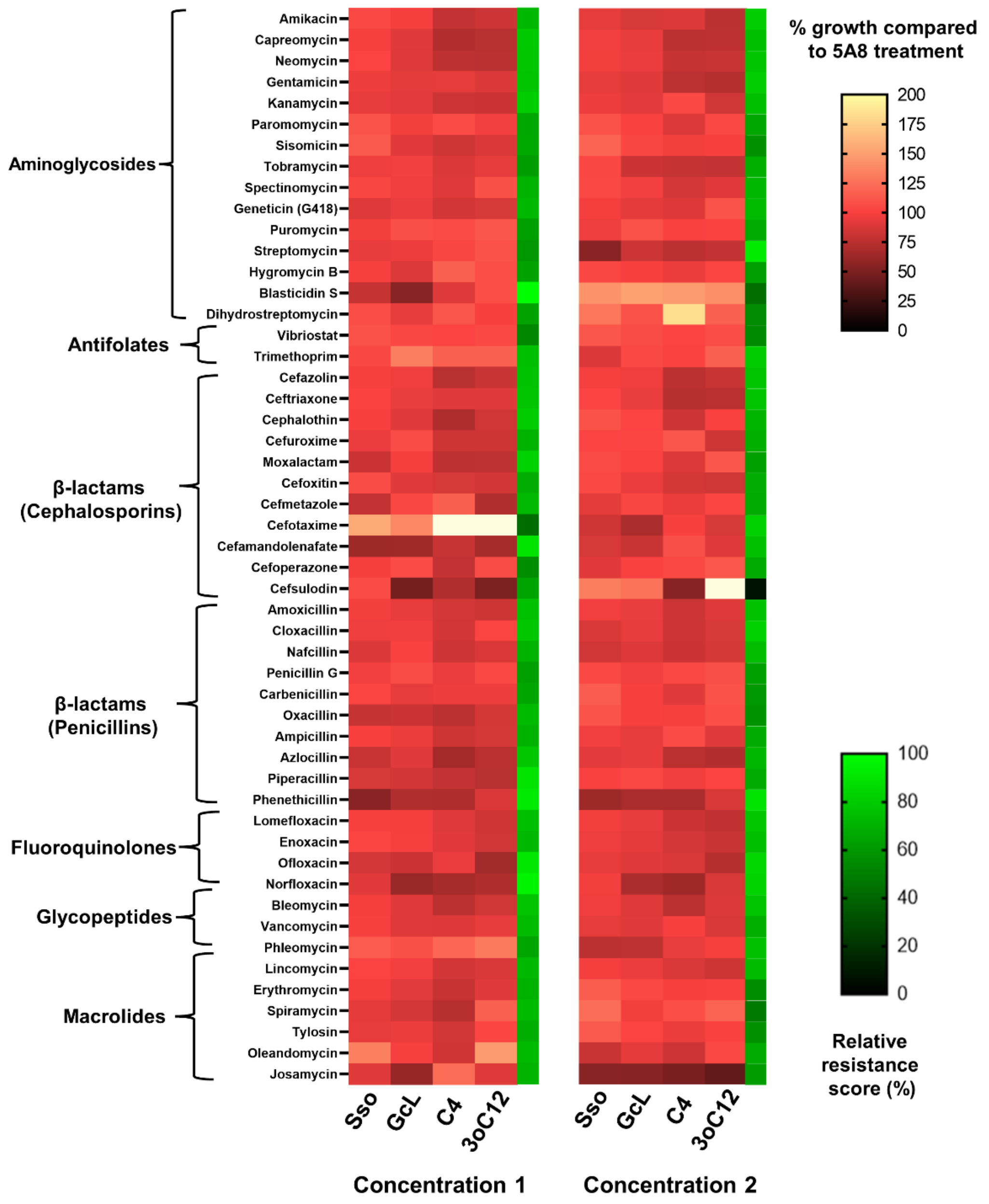


**Fig. S1A: Growth profile of *P. aeruginosa* PA14 with antimicrobials and interference in quorum sensing.** Tested antibiotics and antibacterial compounds (grouped according to their classes and indicated on the left) are at two proprietary dosages. “Concentration 2” is higher than “Concentration 1” (see Methods). Growth of PA14 in the presence of lactonases – SsoPox W263I (Sso) and GcL or AHLs – C4-HSL (C4) and 3-oxo-C12-HSL (3oC12) is represented as a % of PA14 growth in the presence of inactive lactonase SsoPox 5A8 (control) with a white-red-black color scheme. Relative resistance scores (see Methods) are indicated by an additional heat map strip with a green-black color scheme on the right.

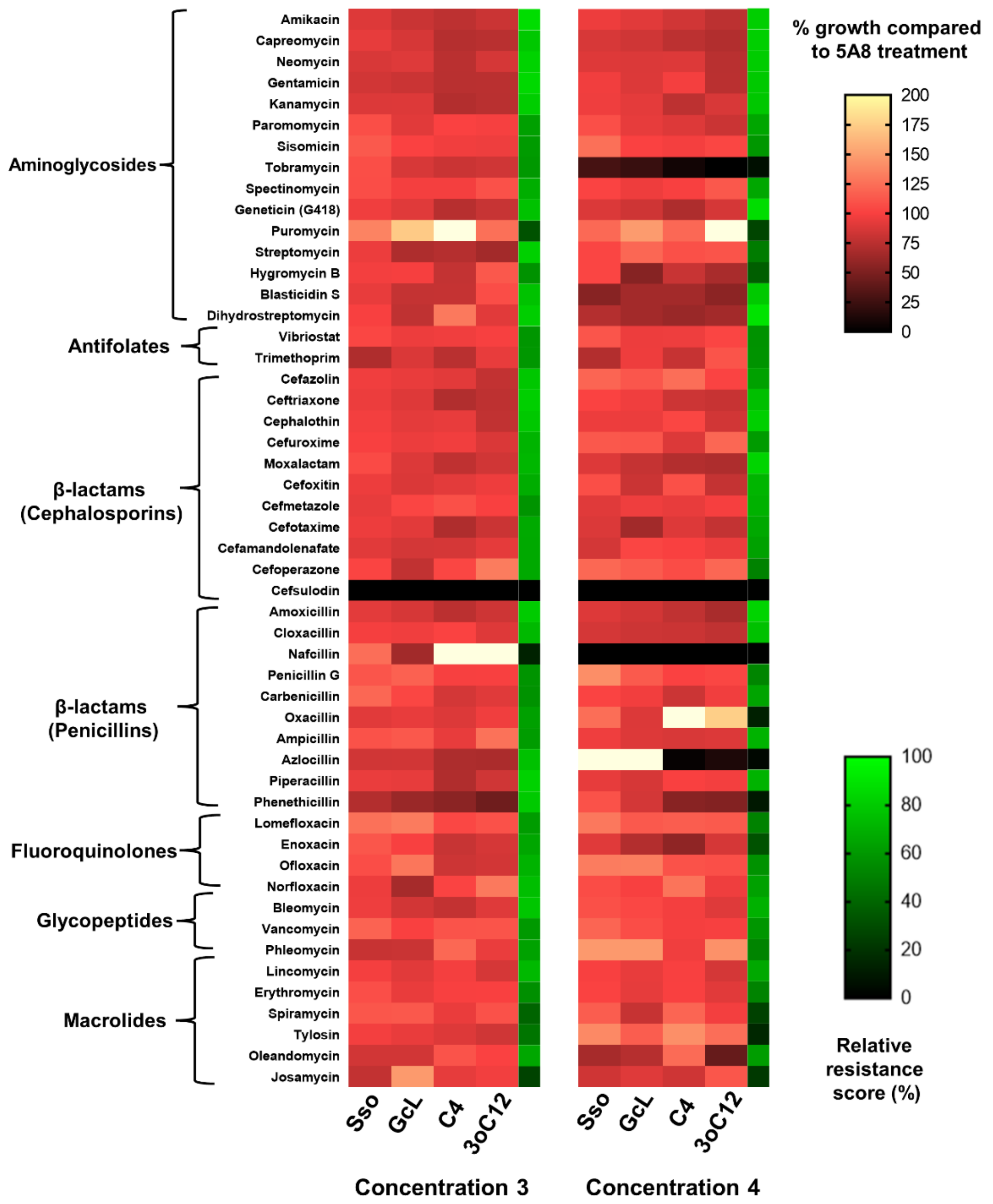


**Fig. S1B: Growth profile of *P. aeruginosa* PA14 with antimicrobials and interference in quorum sensing.** Tested antibiotics and antibacterial compounds (grouped according to their classes and indicated on the left) are at two proprietary dosages. “Concentration 4” is higher than “Concentration 3” (see Methods). Growth of PA14 in the presence of lactonases – SsoPox W263I (Sso) and GcL or AHLs – C4-HSL (C4) and 3-oxo-C12-HSL (3oC12) is represented as a % of PA14 growth in the presence of inactive lactonase SsoPox 5A8 (control) with a white-red-black color scheme. Relative resistance scores (see Methods) are indicated by an additional heat map strip with a green-black color scheme on the right.

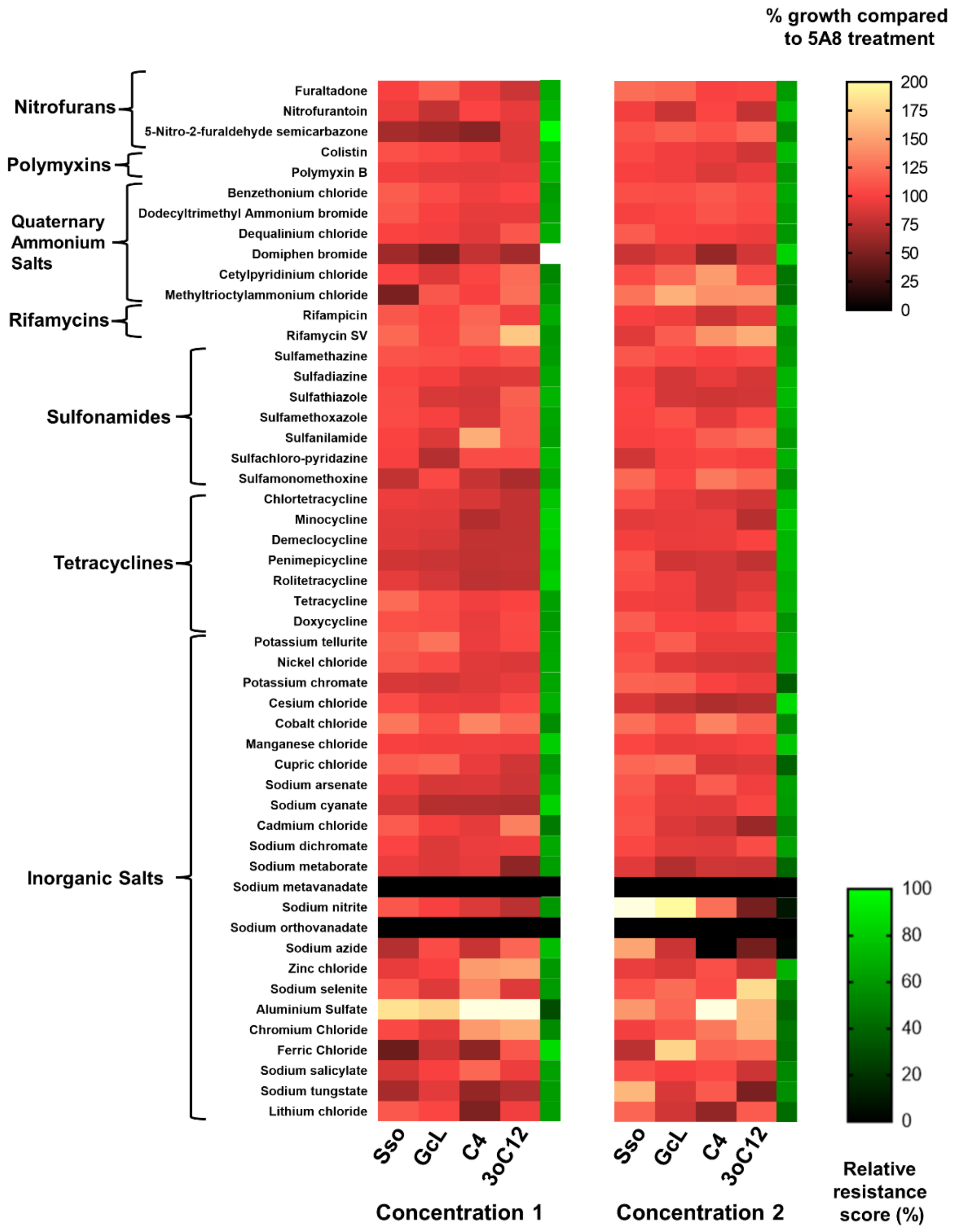


**Fig. S2A: Growth profile of *P. aeruginosa* PA14 with antimicrobials and interference in quorum sensing.** Tested antibiotics and antibacterial compounds (grouped according to their classes and indicated on the left) are at two proprietary dosages. "Concentration 2" is higher than "Concentration 1" (see Methods). Growth of PA14 in the presence of lactonases – SsoPox W263I (Sso) and GcL or AHLs – C4-HSL (C4) and 3-oxo-C12-HSL (3oC12) is represented as a % of PA14 growth in the presence of inactive lactonase SsoPox 5A8 (control) with a white-red-black color scheme. Relative resistance scores (see Methods) are indicated by an additional heat map strip with a green-black color scheme on the right.

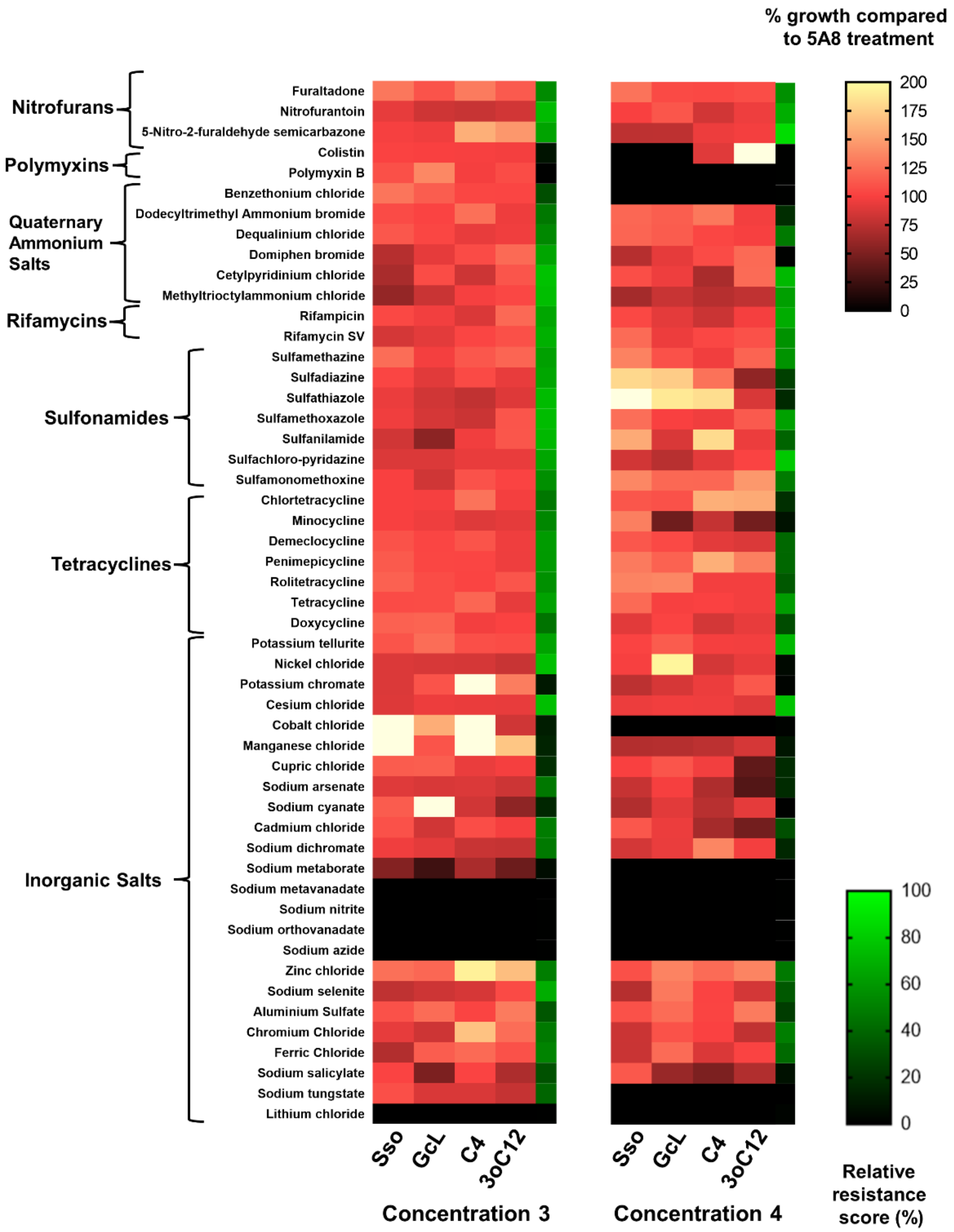


**Fig. S2B: Growth profile of *P. aeruginosa* PA14 with antimicrobials and interference in quorum sensing.** Tested antibiotics and antibacterial compounds (grouped according to their classes and indicated on the left) are at two proprietary dosages. “Concentration 4” is higher than “Concentration 3” (see Methods). Growth of PA14 in the presence of lactonases – SsoPox W263I (Sso) and GcL or AHLs – C4-HSL (C4) and 3-oxo-C12-HSL (3oC12) is represented as a % of PA14 growth in the presence of inactive lactonase SsoPox 5A8 (control) with a white-red-black color scheme. Relative resistance scores (see Methods) are indicated by an additional heat map strip with a green-black color scheme on the right.

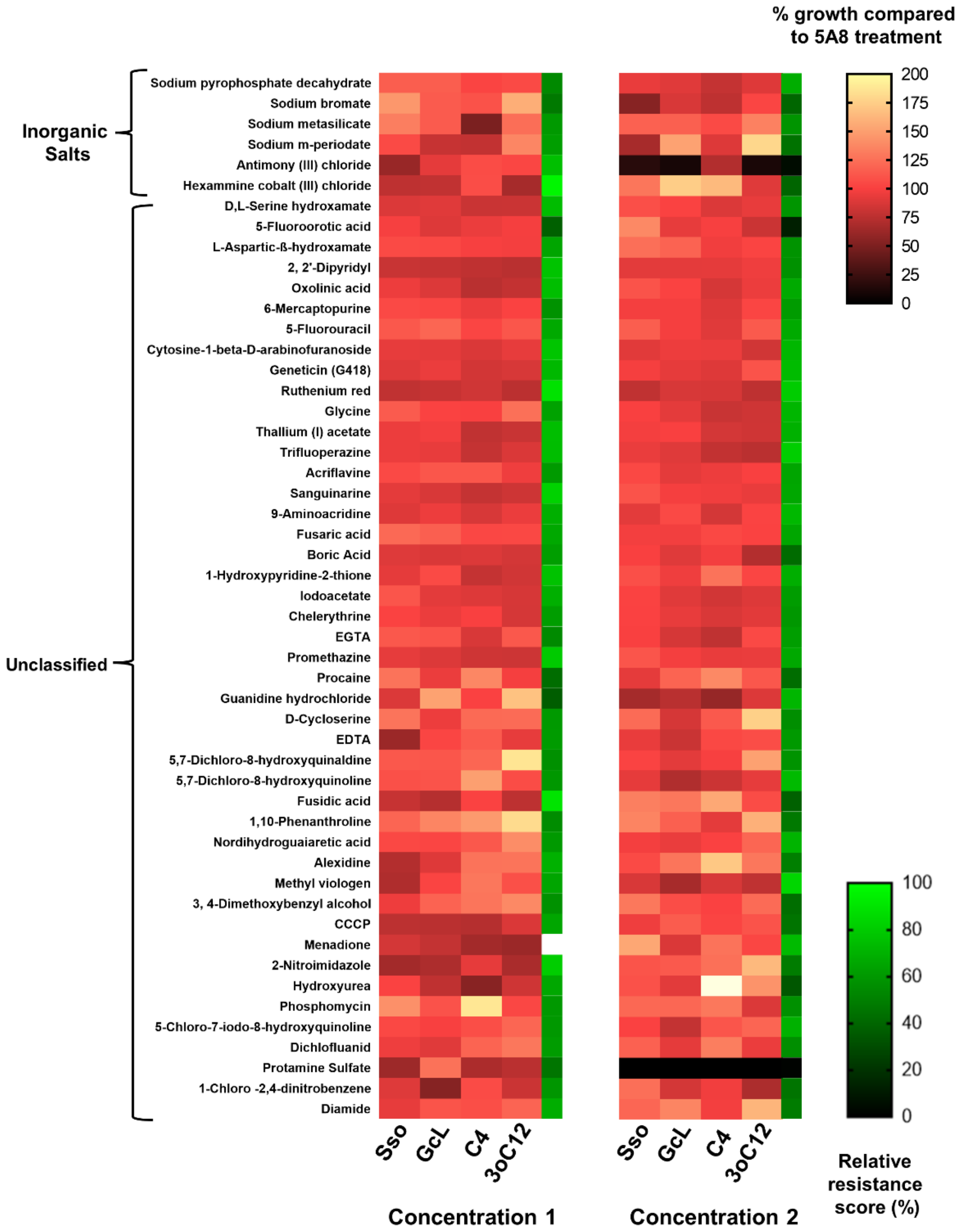


**Fig. S3A: Growth profile of *P. aeruginosa* PA14 with antimicrobials and interference in quorum sensing.** Tested antibiotics and antibacterial compounds (grouped according to their classes and indicated on the left) are at two proprietary dosages. “Concentration 2” is higher than “Concentration 1” (see Methods). Growth of PA14 in the presence of lactonases – SsoPox W263I (Sso) and GcL or AHLs – C4-HSL (C4) and 3-oxo-C12-HSL (3oC12) is represented as a % of PA14 growth in the presence of inactive lactonase SsoPox 5A8 (control) with a white-red-black color scheme. Relative resistance scores (see Methods) are indicated by an additional heat map strip with a green-black color scheme on the right.

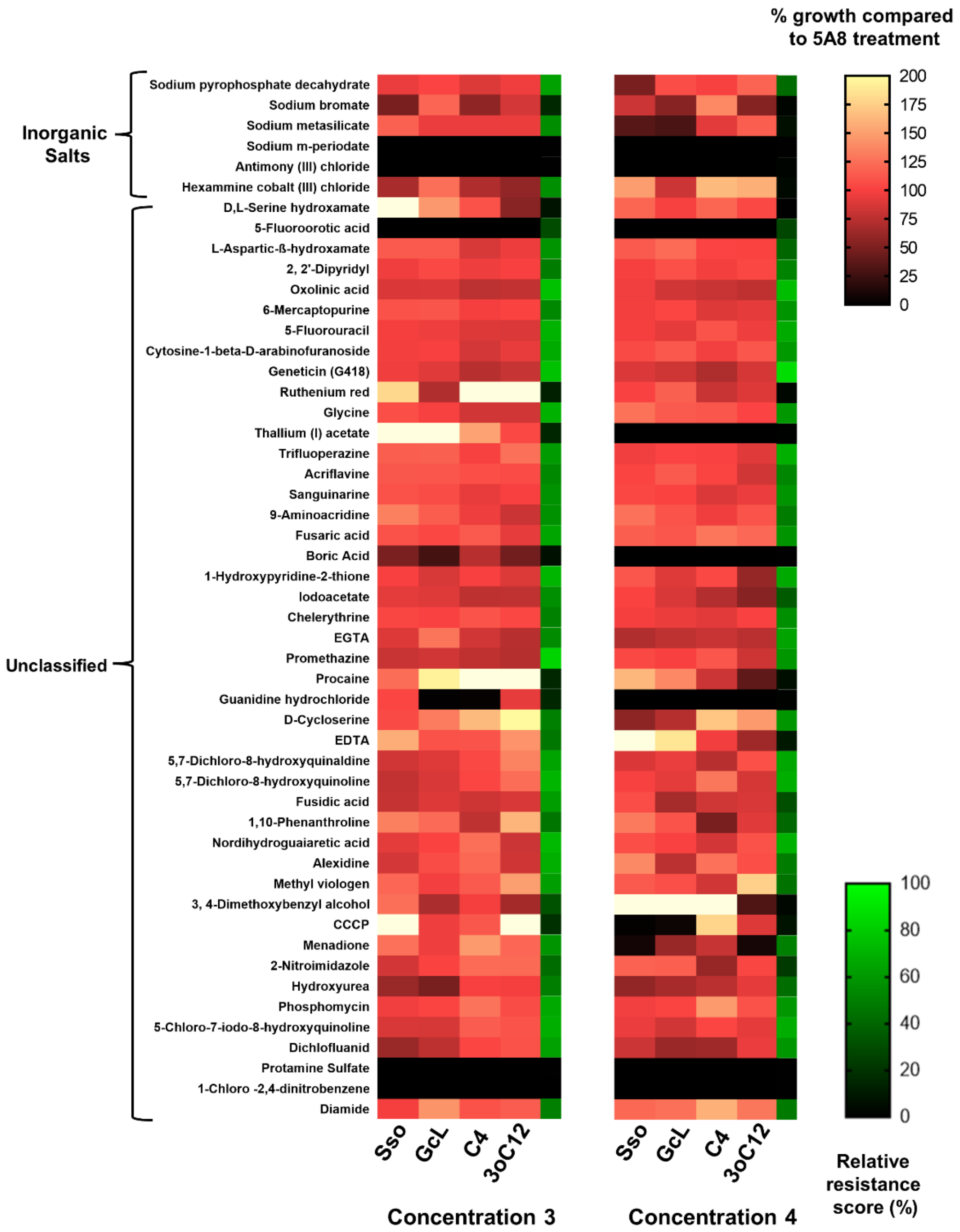


**Fig. S3B: Growth profile of *P. aeruginosa* PA14 with antimicrobials and interference in quorum sensing.** Tested antibiotics and antibacterial compounds (grouped according to their classes and indicated on the left) are at two proprietary dosages. "Concentration 4" is higher than "Concentration 3" (see Methods). Growth of PA14 in the presence of lactonases – SsoPox W263I (Sso) and GcL or AHLs – C4-HSL (C4) and 3-oxo-C12-HSL (3oC12) is represented as a % of PA14 growth in the presence of inactive lactonase SsoPox 5A8 (control) with a white-red-black color scheme. Relative resistance scores (see Methods) are indicated by an additional heat map strip with a green-black color scheme on the right.

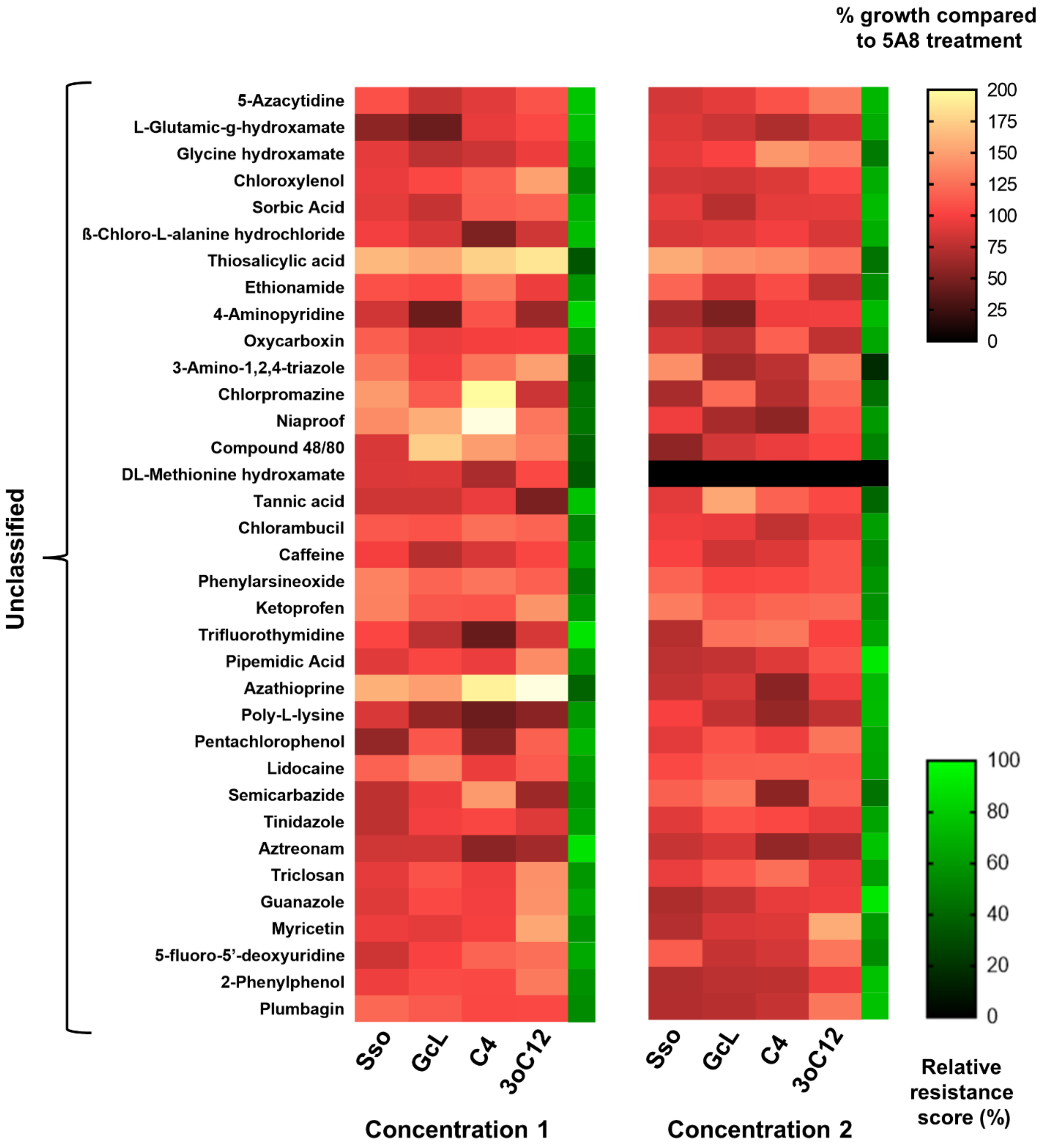


**Fig. S4A: Growth profile of *P. aeruginosa* PA14 with antimicrobials and interference in quorum sensing.** Tested antibiotics and antibacterial compounds (grouped according to their classes and indicated on the left) at two proprietary dosages. "Concentration 2" is higher than "Concentration 1" (see Methods). Growth of PA14 in the presence of lactonases – SsoPox W263I (Sso) and GcL or AHLs – C4-HSL (C4) and 3-oxo-C12-HSL (3oC12) is represented as a % of PA14 growth in the presence of inactive lactonase SsoPox 5A8 (control) with a white-red-black color scheme. Relative resistance scores (see Methods) are indicated by an additional heat map strip with a green-black color scheme on the right.

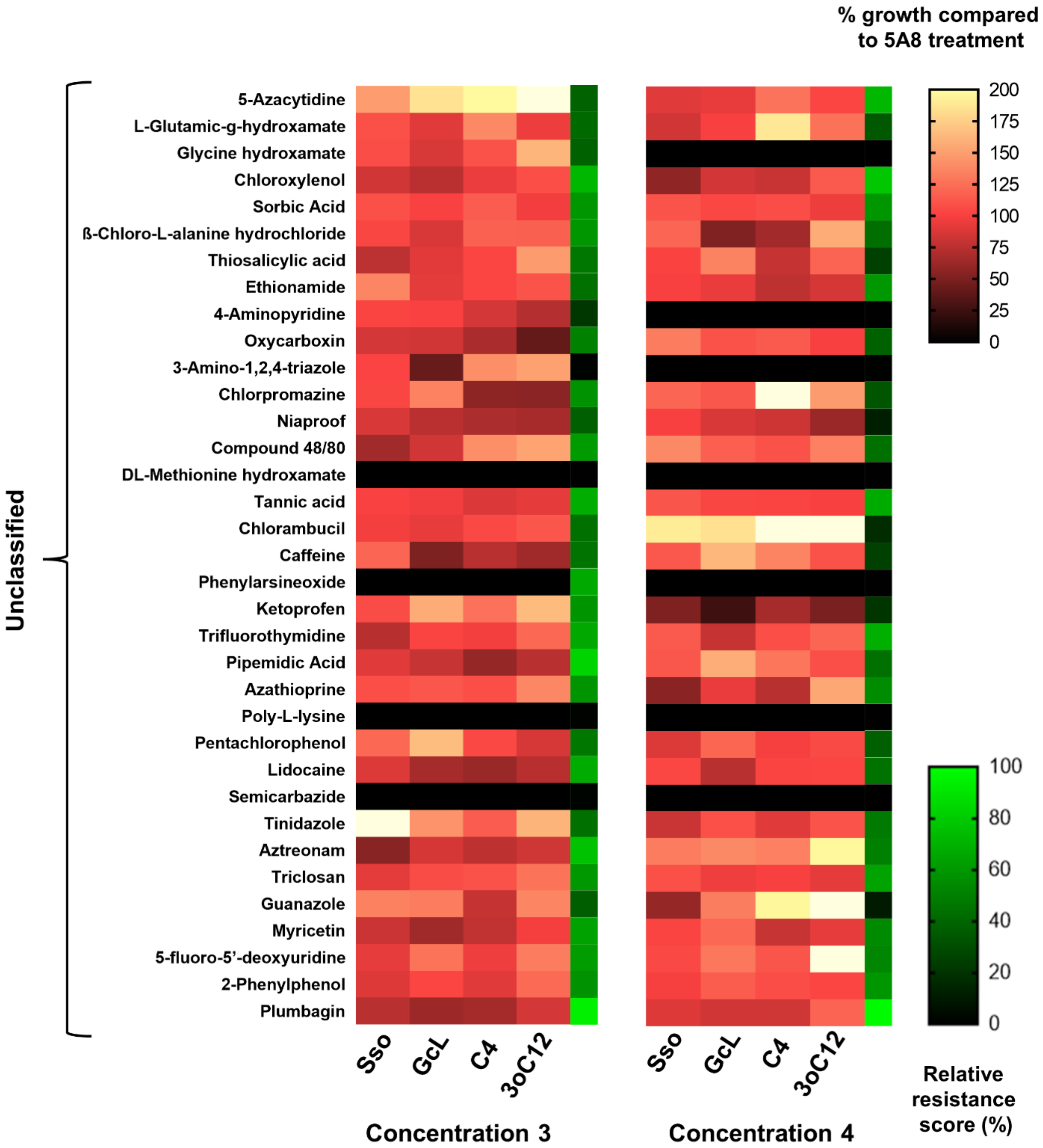


**Fig. S4B: Growth profile of *P. aeruginosa* PA14 with antimicrobials and interference in quorum sensing.** Tested antibiotics and antibacterial compounds (grouped according to their classes and indicated on the left) are at two proprietary dosages. "Concentration 4" is higher than "Concentration 3" (see Methods). Growth of PA14 in the presence of lactonases – SsoPox W263I (Sso) and GcL or AHLs – C4-HSL (C4) and 3-oxo-C12-HSL (3oC12) is represented as a % of PA14 growth in the presence of inactive lactonase SsoPox 5A8 (control) with a white-red-black color scheme. Relative resistance scores (see Methods) are indicated by an additional heat map strip with a green-black color scheme on the right.

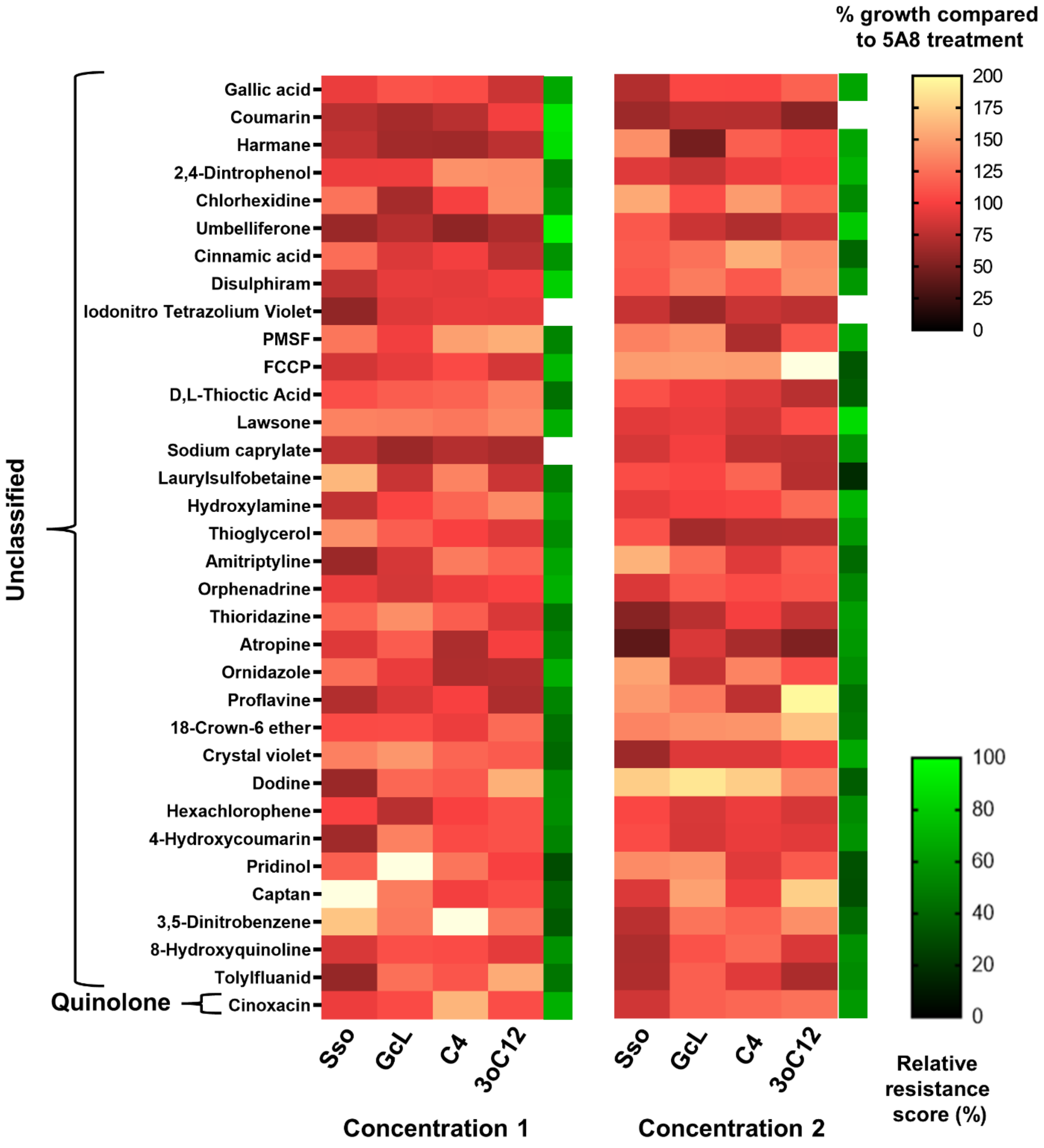


**Fig. S5A: Growth profile of *P. aeruginosa* PA14 with antimicrobials and interference in quorum sensing.** Tested antibiotics and antibacterial compounds (grouped according to their classes and indicated on the left) are at two proprietary dosages. “Concentration 2” is higher than “Concentration 1” (see Methods). Growth of PA14 in the presence of lactonases – SsoPox W263I (Sso) and GcL or AHLs – C4-HSL (C4) and 3-oxo-C12-HSL (3oC12) is represented as a % of PA14 growth in the presence of inactive lactonase SsoPox 5A8 (control) with a white-red-black color scheme. Relative resistance scores (see Methods) are indicated by an additional heat map strip with a green-black color scheme on the right.

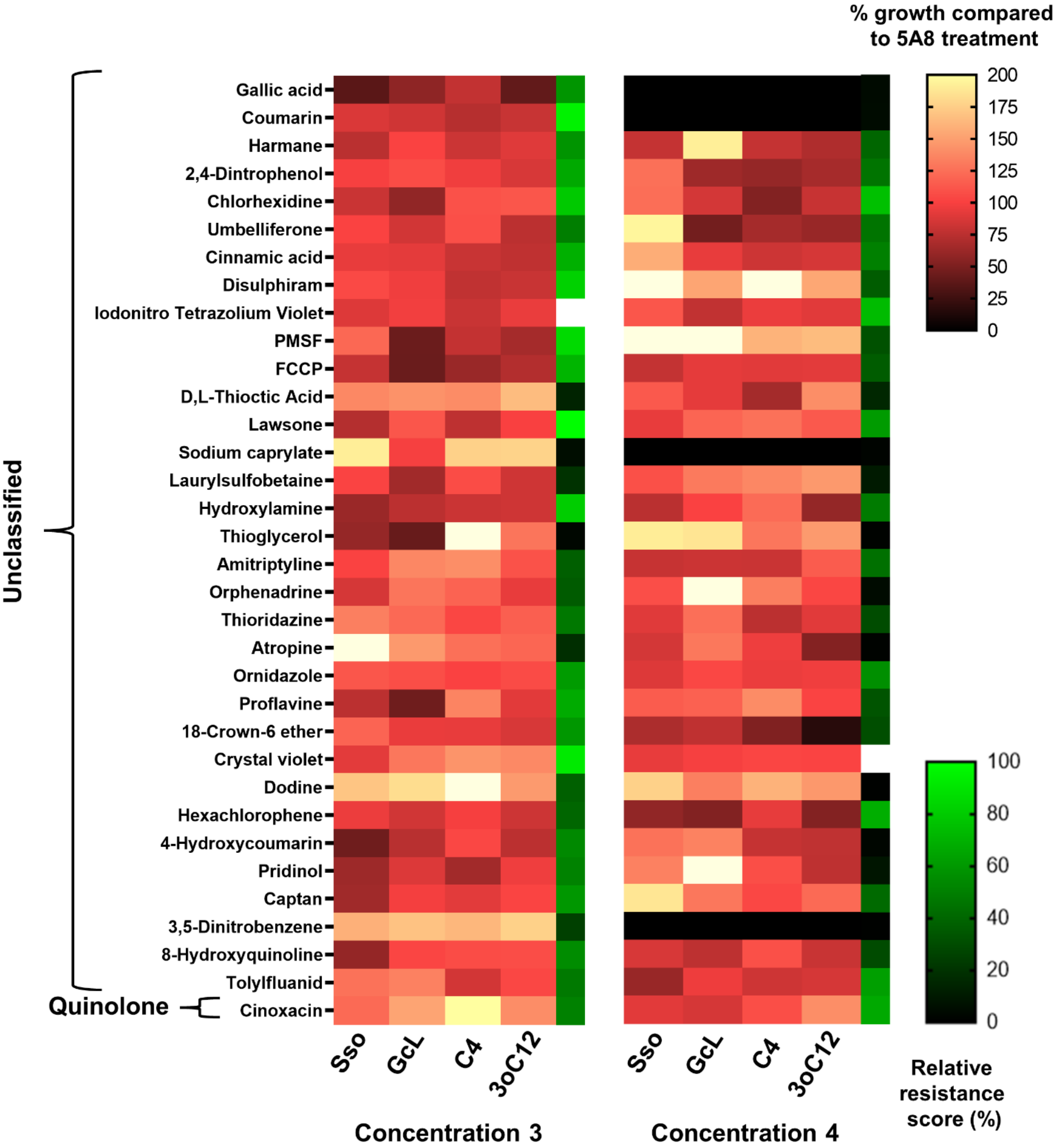


**Fig. S5B: Growth profile of *P. aeruginosa* PA14 with antimicrobials and interference in quorum sensing.** Tested antibiotics and antibacterial compounds (grouped according to their classes and indicated on the left) are at two proprietary dosages. "Concentration 4" is higher than "Concentration 3" (see Methods). Growth of PA14 in the presence of lactonases – SsoPox W263I (Sso) and GcL or AHLs – C4-HSL (C4) and 3-oxo-C12-HSL (3oC12) is represented as a % of PA14 growth in the presence of inactive lactonase SsoPox 5A8 (control) with a white-red-black color scheme. Relative resistance scores (see Methods) are indicated by an additional heat map strip with a green-black color scheme on the right.

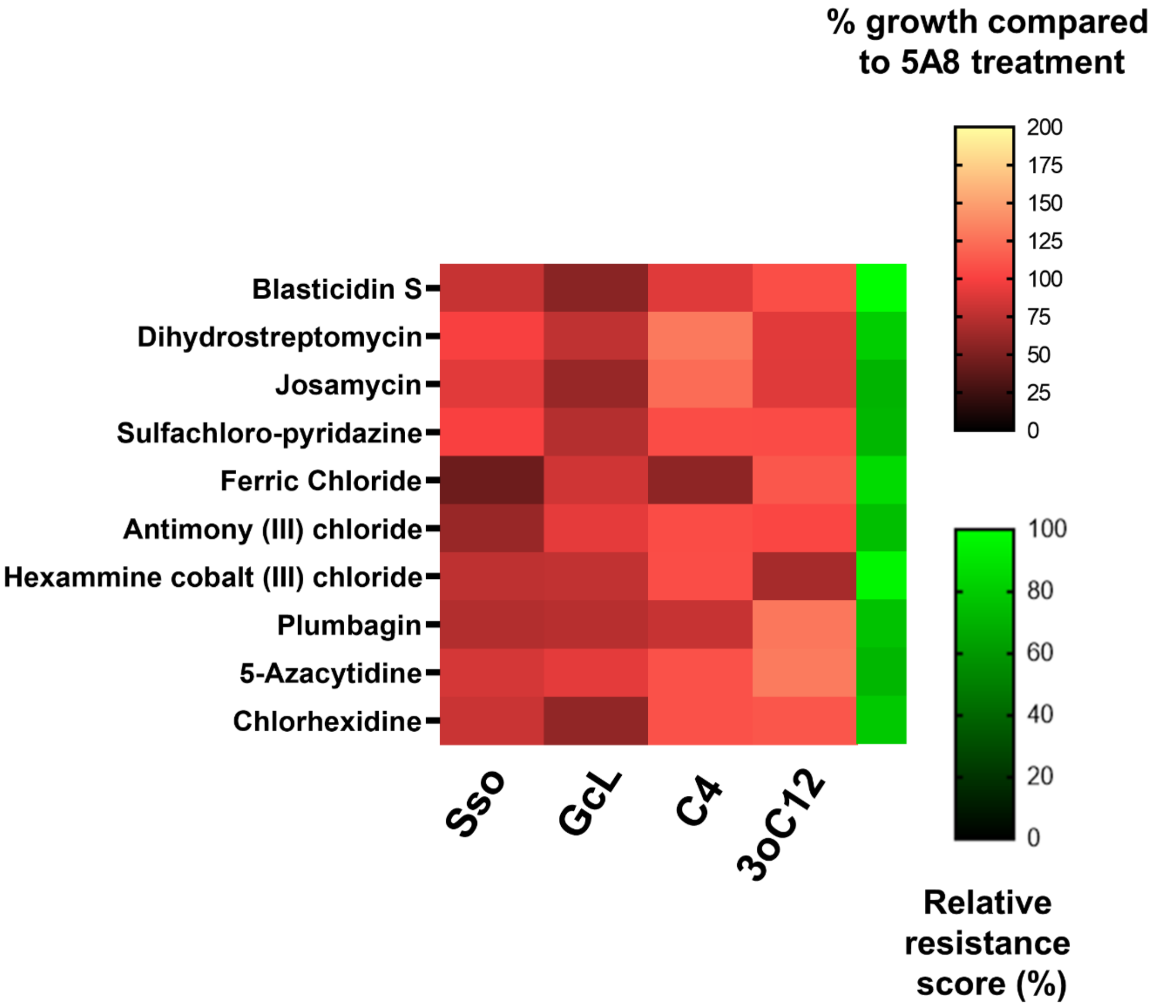


**Fig. S6: Heat maps representing the growth of *P. aeruginosa* PA14 in presence of antibacterial compounds for which the sensitivity of PA14 is regulated by AHL signaling (see main text)**. All these compounds have a relative resistance score of 70 or higher but demonstrate a contrasting change (>25% in at least one treatment) in PA14 growth upon treatment with QQ lactonases or AHLs. Growth of PA14 in the presence of lactonases – SsoPox W263I (Sso) and GcL or AHLs – C4-HSL (C4) and 3-oxo-C12-HSL (3oC12) is represented as a % of PA14 growth in the presence of inactive lactonase SsoPox 5A8 (control) with a white-red-black color scheme. Relative resistance scores (see Methods) are indicated by an additional heat map strip with a green-black color scheme on the right.

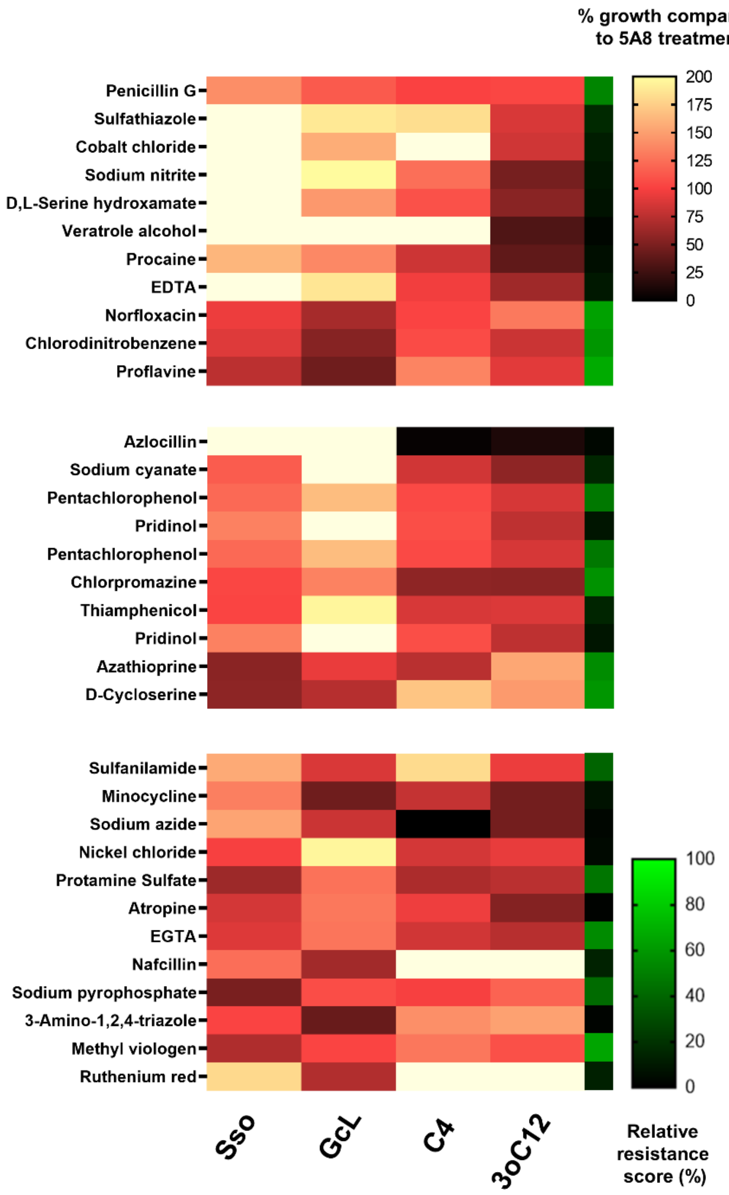


**Fig. S7: Modulations of the antibiotic resistance profile of *P. aeruginosa* PA14 depend on the AHL substrate specificity of QQ lactonase**. Heat maps representing the growth of PA14 in Biolog Phenotype MicroArrays containing indicated antibiotics and antibacterial compounds, for which the growth of PA14 is altered differently by QQ lactonases SsoPox W263I and GcL with different AHL substrate specificities (>25% for at least one lactonase, compared to control). Growth of PA14 in the presence of lactonases – SsoPox W263I (Sso) and GcL or AHLs – C4-HSL (C4) and 3-oxo-C12-HSL (3oC12) is represented as a % of PA14 growth in the presence of inactive lactonase SsoPox 5A8 (control) with a white-red-black color scheme. Relative resistance scores (see Methods) are indicated by an additional heat map strip with a green-black color scheme on the right. **Top heat map panel:** PA14 growth in SsoPox W263I treatment is > GcL. **Middle heat map panel:** PA14 growth in GcL treatment is > SsoPox W263I. **Bottom heat map panel:** contrasting PA14 growth between SsoPox W263I and GcL treatments.

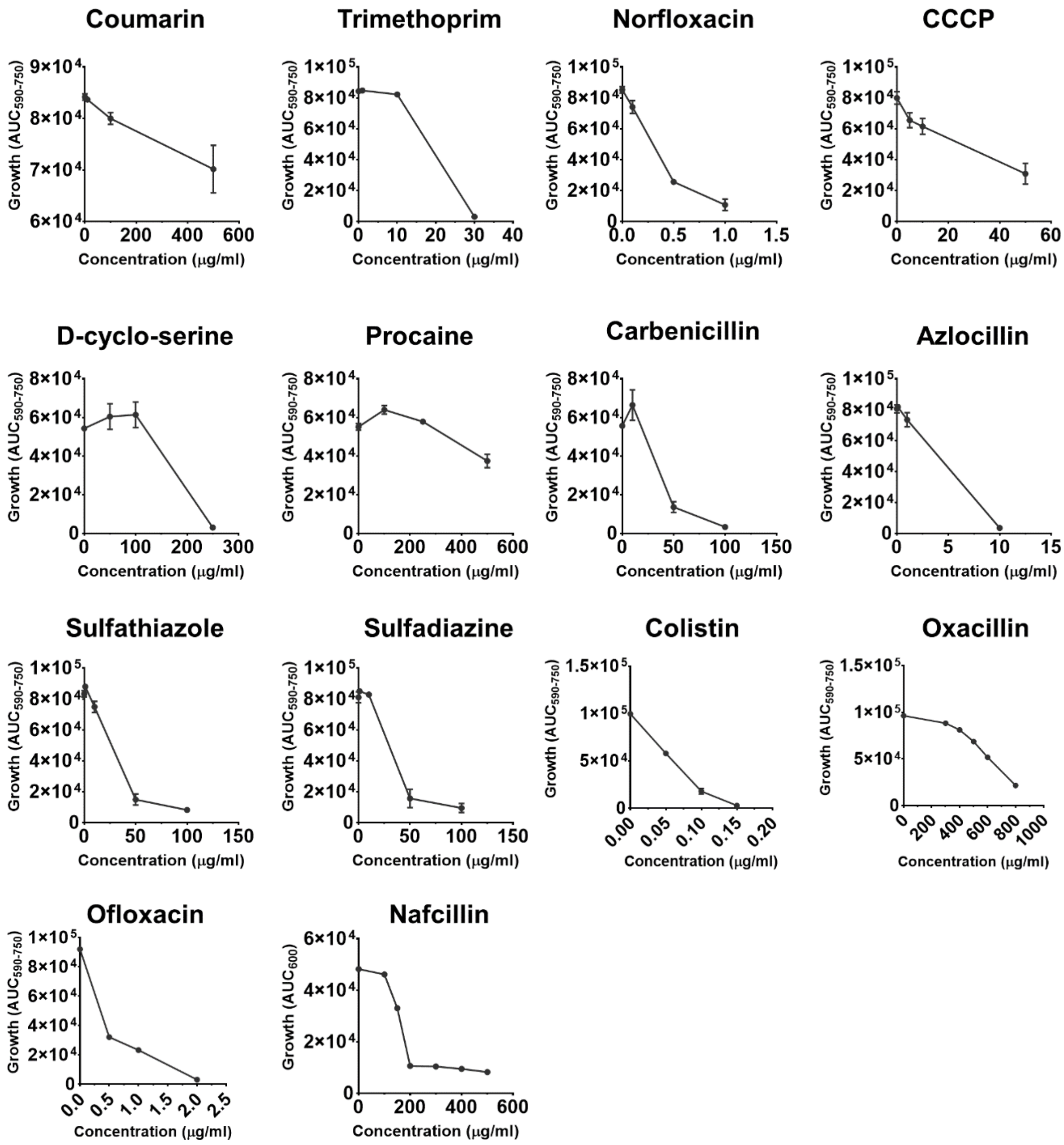


**Fig. S8: Dose-response growth of *P. aeruginosa* PA14 to determine the sublethal testing concentration for several antibiotics and antibacterial compounds**. PA14 was grown in the presence of these compounds at indicated concentrations in Biolog IF-10a media, except in the case of nafcillin, where PA14 was grown in MOPS defined media with 0.2% glucose (Neidhardt *et. al.* 1974). All compounds except for oxacillin, ofloxacin and nafcillin at all concentrations were tested in at least duplicates. Except for nafcillin, where total growth of PA14 was determined by Area-Under-the-Curve (AUC) of $OD_{600}$ growth curve, total growth of PA14 was determined by AUC of the $OD_{590\text{-}750}$ growth curves like Biolog PM experiments.

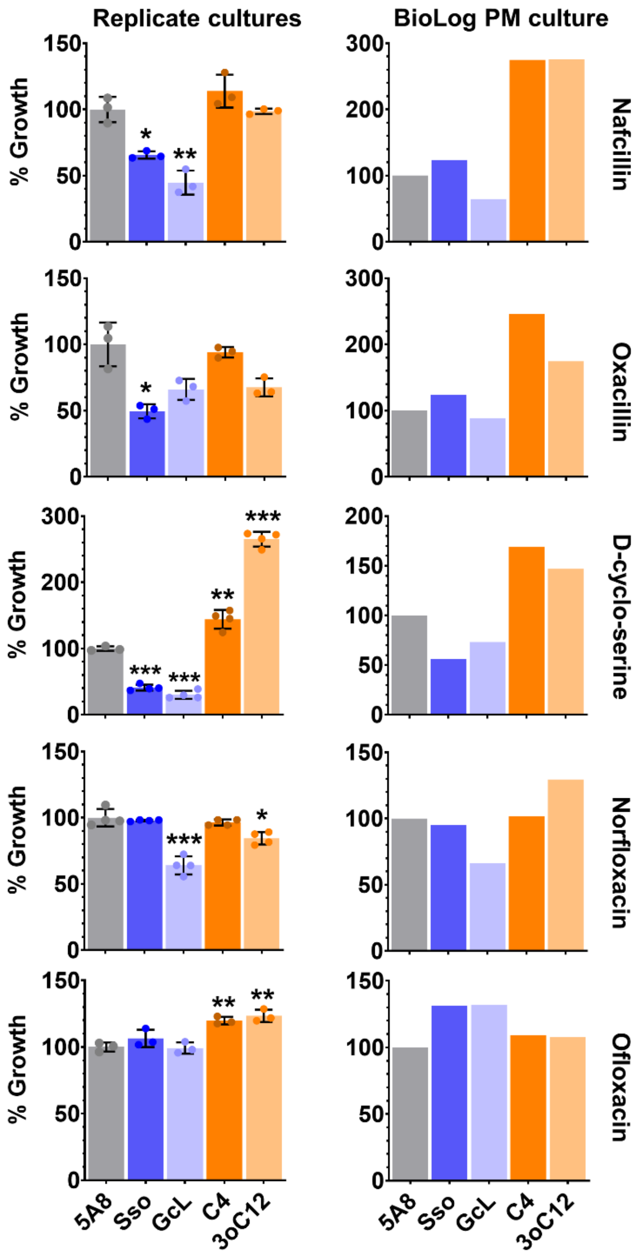


**Fig S9: Comparison between *P. aeruginosa* PA14 growth in Biolog Phenotype MicroArray (PM) cultures and replicated experiments** with 200 µg/mL nafcillin, 800 µg/mL oxacillin, 200 µg/mL D-cyclo-serine, 0.25 µg/mL norfloxacin and 0.5 µg/mL ofloxacin upon treatment with QQ lactonases – SsoPox W263I (Sso) and GcL or AHLs – C4-HSL (C4) and 3-oxo-C12-HSL (3oC12). PA14 growth in all treatments is normalized to their respective 5A8 control (treatment with inactive lactonase SsoPox 5A8) set at 100%. All lactonases and AHLs are used at 50 µg/mL and 10 µM final concentrations, respectively. All experiments were done, and all data is represented as the mean and standard deviation of at least triplicates, for the replicated cultures. Statistical significance of all treatments compared to the control (5A8) was calculated using unpaired two-tailed t-tests with Welch's correction and significance values are indicated as - ***$p < 0.0005$, **$p < 0.005$ and *$p < 0.05$.

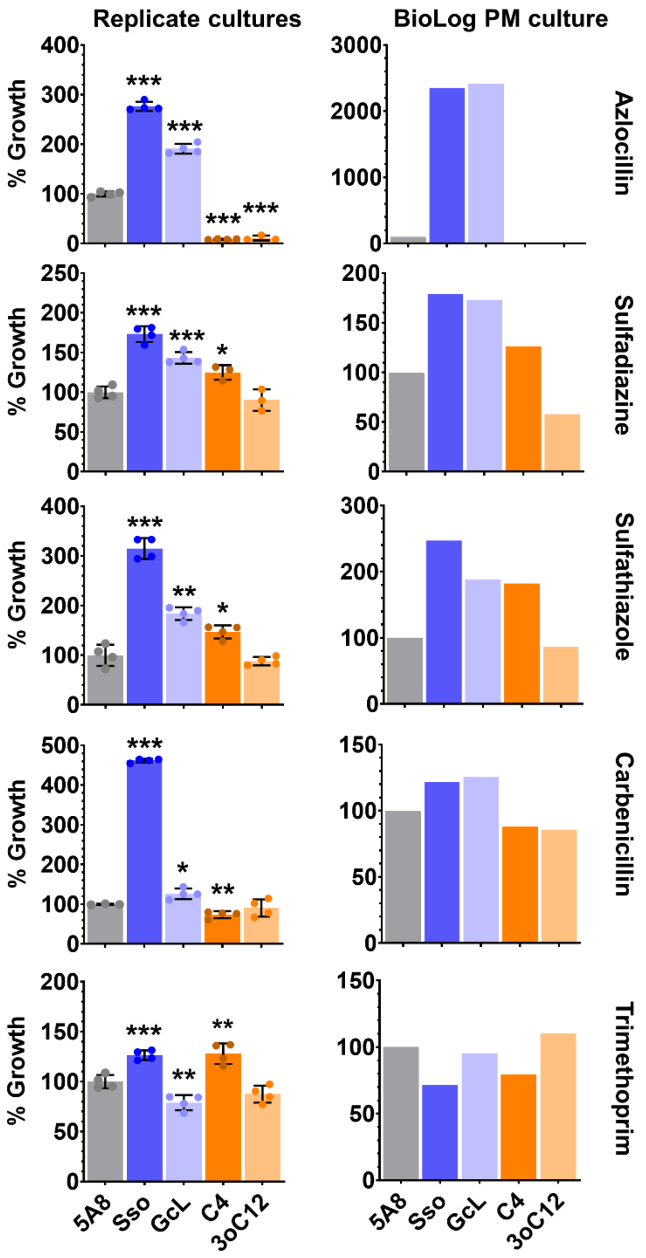


**Fig S10: Comparison between *P. aeruginosa* PA14 growth in Biolog Phenotype MicroArray (PM) cultures and replicated experiments** with 5 µg/mL azlocillin, 25 µg/mL sulfadiazine, 25 µg/mL sulfathiazole, 50 µg/mL carbenicillin and 20 µg/mL trimethoprim upon treatment with QQ lactonases – SsoPox W263I (Sso) and GcL or AHLs – C4-HSL (C4) and 3-oxo-C12-HSL (3oC12). PA14 growth in all treatments is normalized to their respective 5A8 control (treatment with inactive lactonase SsoPox 5A8) set at 100%. All lactonases and AHLs are used at 50 µg/mL and 10 µM final concentrations, respectively. All experiments were done, and all data is represented as the mean and standard deviation of at least triplicates, for the replicated cultures. Statistical significance of all treatments compared to the control (5A8) was calculated using unpaired two-tailed t-tests with Welch's correction and significance values are indicated as - ***$p < 0.0005$, **$p < 0.005$ and *$p < 0.05$.

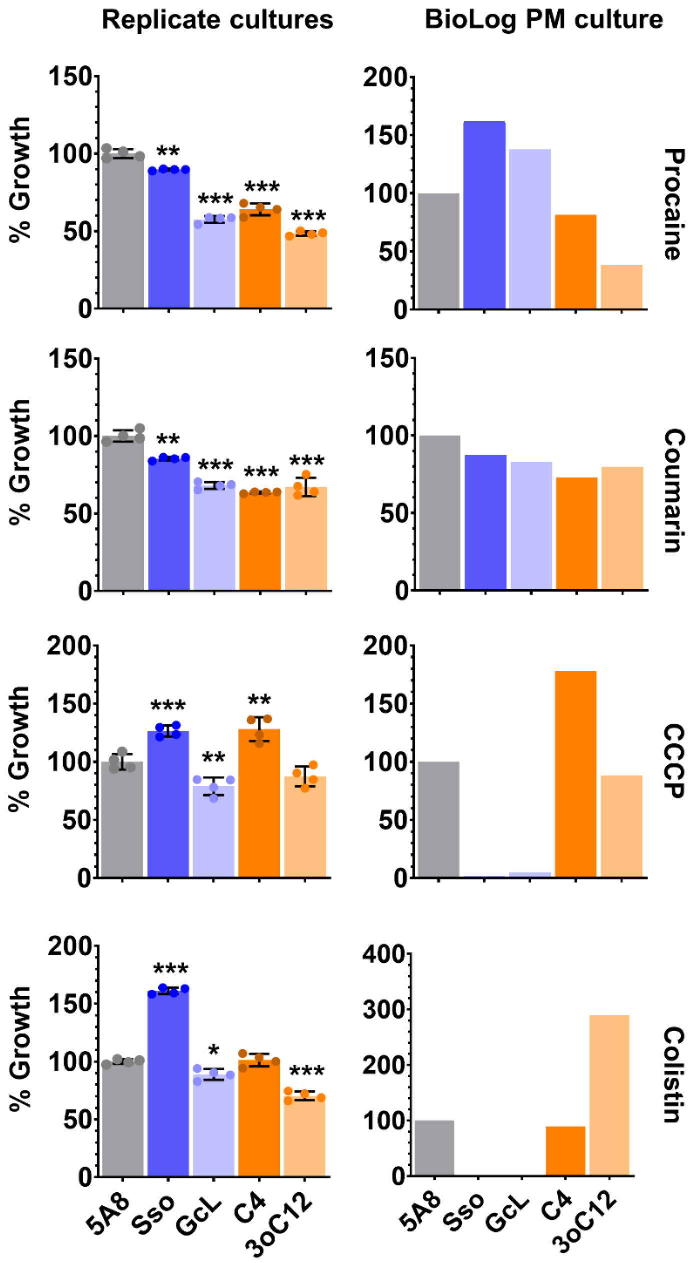


**Fig. S11: Comparison between *P. aeruginosa* PA14 growth in Biolog Phenotype MicroArray (PM) cultures and replicated experiments** with 500 µg/mL procaine, 500 µg/mL coumarin, 50 µg/mL carbonyl cyanide 3-chlorophenylhydrazone or CCCP and 0.1 µg/mL colistin upon treatment with QQ lactonases – SsoPox W263I (Sso) and GcL or AHLs – C4-HSL (C4) and 3-oxo-C12-HSL (3oC12). PA14 growth in all treatments is normalized to their respective 5A8 control (treatment with inactive lactonase SsoPox 5A8) set at 100%. All lactonases and AHLs are used at 50 µg/mL and 10 µM final concentrations, respectively. All experiments were done, and all data is represented as the mean and standard deviation of at least triplicates, for the replicated cultures. Statistical significance of all treatments compared to the control (5A8) was calculated using unpaired two-tailed t-tests with Welch's correction and significance values are indicated as - $***p < 0.0005$, $**p < 0.005$ and $*p < 0.05$.

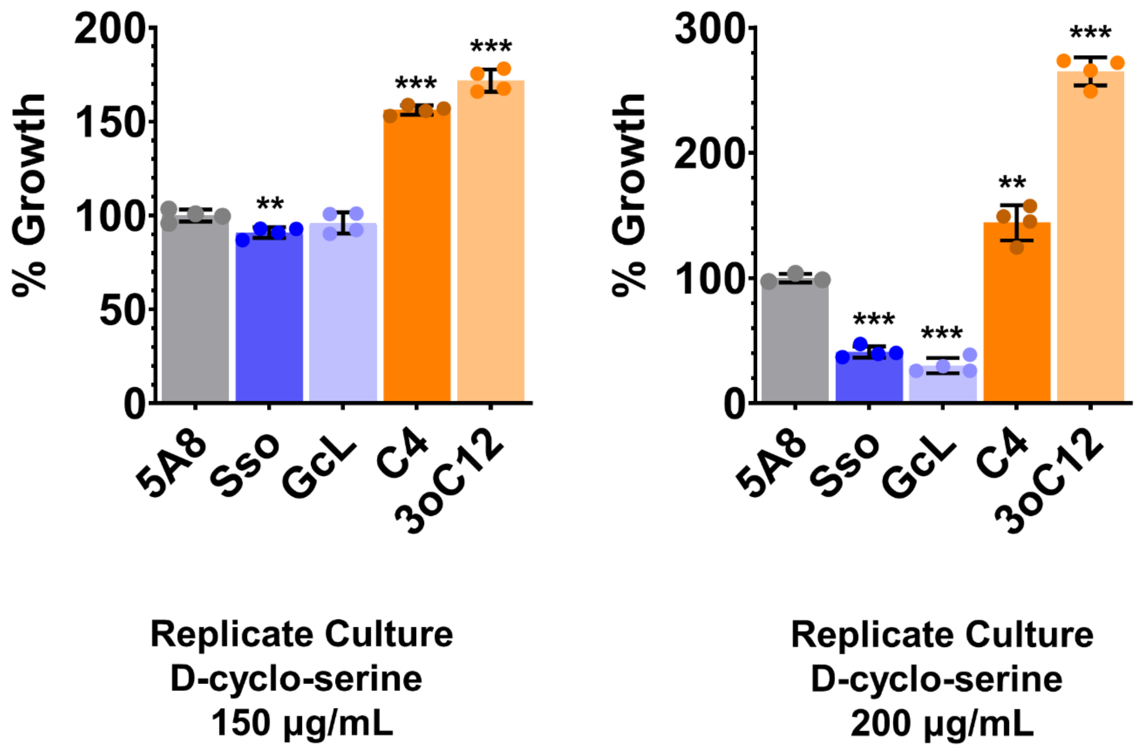


**Fig. S12: *P. aeruginosa* PA14 growth in replicated experiments with antibiotics is sensitive to changes in the sublethal concentration of the antibiotic**. Growth of PA14 in the presence of 150 µg/mL and 200 µg/mL D-cyclo-serine upon treatment with QQ lactonases – SsoPox W263I (Sso) and GcL or AHLs – C4-HSL (C4) and 3-oxo-C12-HSL (3oC12) is shown. PA14 growth in all treatments is normalized to their respective 5A8 control (treatment with inactive lactonase SsoPox 5A8) set at 100%. All lactonases and AHLs are used at 50 µg/mL and 10 µM final concentrations, respectively. All experiments were done, and all data is represented as the mean and standard deviation of at least triplicates, for the replicated cultures. Statistical significance of all treatments compared to the control (5A8) was calculated using unpaired two-tailed t-tests with Welch's correction and significance values are indicated as - ****p < 0.0005*, ***p < 0.005* and **p < 0.05*.

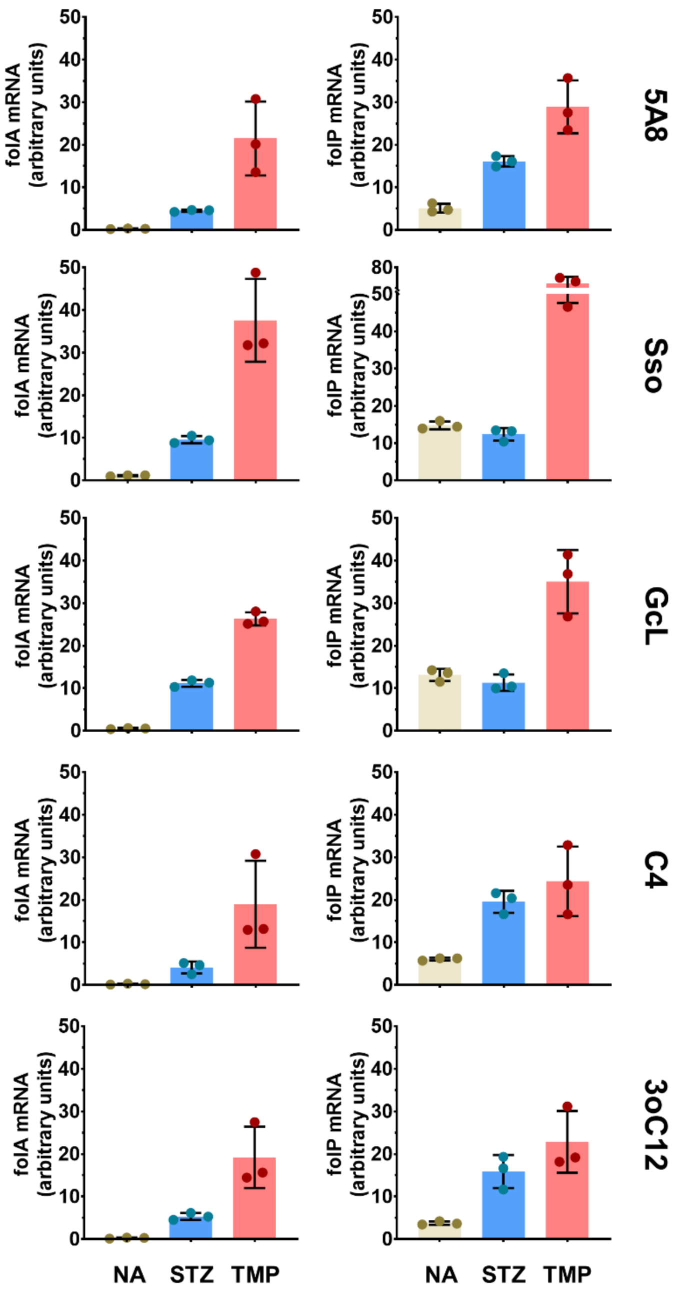


**Fig. S13: qRT-PCR quantifications of changes in transcript mRNA levels of genes folA and folP** in *P. aeruginosa* PA14 without (No Antibiotic, NA) or with antibiotics - 25 µg/mL sulfathiazole (STZ) and 20 µg/mL trimethoprim (TMP) upon treatment with QQ lactonases – SsoPox W263I (Sso) and GcL or AHLs – C4-HSL (C4) and 3-oxo-C12-HSL (3oC12), compared to control treatment (inactive lactonase SsoPox 5A8, indicated as 5A8). folA and folP mRNA levels in all treatments were determined using the relative quantification method using the recA gene as endogenous control and represented using arbitrary units. All lactonases and AHLs are used at 50 µg/mL and 10 µM final concentrations, respectively. All experiments were done, and all data is represented as the mean and standard deviation of at least triplicates.
